# Twitching cells use a chemoreceptor to detect bacterial competitors

**DOI:** 10.1101/2022.11.28.518211

**Authors:** Kaitlin D. Yarrington, Tyler N. Shendruk, Dominique H. Limoli

**Affiliations:** Department of Microbiology and Immunology, Carver College of Medicine, University of Iowa, Iowa City, IA, United States; School of Physics and Astronomy, The University of Edinburgh, Edinburgh EH9 3FD, United Kingdom

## Abstract

Bacteria live in cosmopolitan communities, where the ability to sense and respond to interspecies and environmental signals is critical for survival. We previously showed the pathogen *Pseudomonas aeruginosa* detects secreted peptides from bacterial competitors and navigates interspecies signal gradients using pilus-based motility. Yet, it remained unknown whether *P. aeruginosa* utilizes a designated chemosensory system for this behavior. Here, we performed a comprehensive genetic analysis of a putative pilus chemosensory system to reveal behaviors of mutants that retain motility, but are blind to interspecies signals. The enzymes predicted to methylate (PilK) and demethylate (ChpB) the putative pilus chemoreceptor, PilJ, are necessary for cells to control the direction of migration. While these findings implicate PilJ as a *bona fide* chemoreceptor, such function had yet to be experimentally defined, as PilJ is essential for motility. Thus, we constructed systematic genetic modifications of PilJ and found that without the predicted ligand binding domains or methylation sites cells lose the ability to detect competitor gradients, despite retaining pilus-mediated motility. Collectively, this work uncovers the chemosensory nature of PilJ, providing insight into chemotactic interactions necessary for bacterial survival in polymicrobial communities and revealing putative pathways where therapeutic intervention might disrupt bacterial communication.

## Introduction

Microbes often exist in complex, dynamic environments and have evolved sophisticated systems to perceive and respond to the outside world. Because they commonly reside in multispecies communities, bacteria experience gradients of nutrients, metabolites, and secretions generated by neighboring cells. Gradients are particularly steep in surface-attached biofilm communities and ecological theory predicts bacteria must sense and respond to competitor and cooperator signals to thrive in such complex environments (Foster & Bell, 2012; Oliveira et al., 2016).

In line with this hypothesis, we recently reported that *Pseudomonas aeruginosa* is attracted to gradients of secreted factors from other microbial species (Limoli et al., 2019). *Pseudomonads* are opportunistic bacteria found in polymicrobial communities in soil, wounds, and chronic lung infections, such as those in people with cystic fibrosis (Limoli & Hoffman, 2019; Tashiro et al., 2013). *P. aeruginosa* is frequently co-isolated with *Staphylococcus aureus* from cystic fibrosis respiratory samples and coinfections can persist for decades (Deleon et al., 2014; Gabrilska & Rumbaugh, 2015; Hotterbeekx et al., 2017; Limoli & Hoffman, 2019). Coinfection is also associated with pulmonary decline; thus, understanding ecological competition between these organisms may provide insight into patient outcomes (Hubert et al., 2013; Limoli et al., 2016; Maliniak et al., 2016).

Accordingly, *in vitro* studies have documented interspecies interactions between *P. aeruginosa* and *S. aureus* leading to reciprocal enhancement of antibiotic tolerance, production of virulence factors, and ability to alter host immune cell responses, further supporting clinical observations (Limoli & Hoffman, 2019; Orazi et al., 2020; Orazi & O’Toole, 2017; Orazi et al., 2019). Additionally, these data suggest each species may sense a secreted signal from the other, which instigates a competitive or cooperative response through alteration of their virulence arsenals, a model supported by differential regulation of specific *P. aeruginosa* virulence pathways in response to *S. aureus* exoproducts (Kvich et al., 2022; Zarrella & Khare, 2022). Remarkably, *P. aeruginosa* and *S. aureus* have been shown to form mixed microcolonies when cocultured on bronchial epithelial cells (Orazi & O’Toole, 2017). One possible explanation for the formation of mixed communities is that *P. aeruginosa* and *S. aureus* may be initially attracted to one another through detection of secreted interspecies signals. Such attraction has the potential to facilitate formation of blended microcolonies or microbial competition, depending on the environmental conditions.

Supporting this model, *P. aeruginosa* senses secreted *Staphylococcal* peptide toxins referred to as phenol soluble modulins (PSMs) and responds with directed motility towards the increasing PSM concentration gradient, mediated by the type IV pilus (TFP) (Limoli et al., 2019). With the identification of a putative interspecies chemoattractant for pilus-based motility, we hypothesized that *P. aeruginosa* uses a chemosensory pathway to move towards *S. aureus*. TFP-mediated motility, or twitching motility, occurs through the grappling hook activity of the pilus, which undergoes episodes of extension, substrate attachment and retraction which pulls the cell body along the surface (Burrows, 2012). The direction of twitching motility is thought to be controlled by preferential extension of pili at the pole facing the direction of movement, referred to as the leading pole. Cells are predicted to change direction by extending pili from the opposite pole, reversing the direction of cellular movement along the long axis of the cell body and so swapping leading poles (Kühn et al., 2021). However, whether modulation of reversal frequency is necessary and sufficient to bias the movement of twitching cells towards a chemoattractant, similar to the run-and-tumble or run-reverse-turn strategies used in flagella-mediated chemotaxis, remains unknown (Qian et al., 2013). While planktonic swimming cells smoothly sample gentle chemoattractant gradients by typically traveling a full body length or more between tumbling events, surface-associated twitching motility is hundreds to thousands of times slower and steep, varying gradients characterize the chemotactic landscape (Berg & Brown, 1972; Patteson et al., 2015). Thus, twitching cells experience less certain chemotactic signals (Carabelli et al., 2020; Hook et al., 2019; Oliveira et al., 2016). Thus, we predict that additional parameters need to be considered to fully understand how a twitching community biases directional movement. Indeed, even simple models of twitching motility are known to produce complex dynamics (Nagel et al., 2020).

While TFP-mediated chemotaxis has not been thoroughly dissected in *P. aeruginosa*, prior work has described chemotactic roles for the two proteins of the putative pilus chemosensory system, Pil-Chp (Kühn et al., 2021; Oliveira et al., 2016). Namely, the predicted response regulators, PilG and PilH, are thought to control reversals and increase levels of the intracellular second messenger cyclic adenosine monophosphate (cAMP) through PilG direct activation of the adenylate cyclase CyaB (Fulcher et al., 2010; Kühn et al., 2021; Oliveira et al., 2016; Persat et al., 2015). cAMP controls a large arsenal of virulence factors targeting both eukaryotic and prokaryotic cells, as well as multiple modes of motility through activation of the virulence response transcriptional regulator (Vfr) (Wolfgang et al., 2003). However, whether cAMP is also necessary to transduce the detection of interspecies attractants to modulate directional motility has not been investigated.

In addition, Pil-Chp includes homologous proteins to a majority of the CheI flagella chemotaxis system in *P. aeruginosa* (Matilla et al., 2021; Sampedro et al., 2015). However, unlike the 24 CheI-associated chemoreceptors, referred to as methyl-accepting chemotaxis proteins (MCPs), Pil-Chp only has one MCP, called PilJ. This putative pilus chemoreceptor differs from the flagella systems, in that PilJ is uniquely essential for twitching motility and possesses low protein sequence similarity in both the ligand binding domain (LBD) and cytoplasmic signaling domains with other MCPs (Delange et al., 2007; Matilla et al., 2021). The chemoreceptor for this system and the chemotactic role of the rest of the pathway has yet to be fully interrogated.

To determine the contribution of the remaining proteins encoded within the Pil-Chp pathway, we systematically deleted these genes and identified those mutants that retain twitching yet are unable to bias movement up a gradient of *S. aureus* secreted factors. From this analysis, six genes fit these criteria: *pilK*, *chpB*, *chpC*, *pilH*, *cyaB*, and *cpdA*. ChpC is a CheW-like linker protein and PilH is a response regulator, while CyaB and CpdA are enzymes that synthesize and degrade cAMP, respectively. PilK and ChpB are proteins that control chemoreceptor adaptation through methylation of the Pil-Chp MCP PilJ, implicating a role for PilJ as a chemoreceptor for interspecies signals. Accordingly, PilJ mutations in the key regions that define an MCP, including the LBD for sensing interspecies signals and methylation sites for chemoreceptor adaptation, revealed PilJ is necessary for *P. aeruginosa* to perceive and bias movements towards *S. aureus.* Quantification of cAMP in single cells also reveals that cAMP levels rise in *P. aeruginosa* during chemotaxis towards *S. aureus*; yet cAMP increases are not strictly due to enhanced twitching motility. Collectively, these results define a novel chemosensory role for PilJ and the Pil-Chp system to sense and respond to interspecies signals.

## Results: The Pil-Chp system controls *P. aeruginosa* attraction to *S. aureus* peptides

The directional nature of *P. aeruginosa* movement up a gradient of *S. aureus* secreted peptides suggests a role for a chemosensory network. We hypothesize that Pil-Chp controls a chemotaxis-like TFP-mediated response by *P. aeruginosa* towards *S. aureus*. To test this hypothesis, we systematically deleted genes in several components of the Pil-Chp pathway to identify mutants that retain twitching motility, but show diminished directional response to *S. aureus*, using a macroscopic directional motility assay (Figure 1A, B) (Kearns et al., 2001; Limoli et al., 2019). Growth medium or cell-free *S. aureus* supernatant was spotted on top of an agar plate and allowed to diffuse for 24 hours to form a gradient. *P. aeruginosa* was then spotted at a distance from the gradient and allowed to respond to each gradient for 36 hours before imaging the plates and quantifying the directional motility ratio (Figure 1C). Four of the 16 pilus mutants retain twitching motility, but reduced directional motility up a gradient of *S. aureus* secreted signals (Figure 1D, E). The remaining mutants either phenocopy wildtype or are non-motile (Figure 1A,B). Mutants with reduced ability to respond include genes that encode for the predicted methyltransferase PilK and methylesterase ChpB. The other two have modification of the chemoreceptor ChpC, which is a CheW-like linker protein that connects the chemoreceptor and kinase, and PilH, which is a response regulator that is predicted to regulate pilus retraction (Darzins, 1994, 2006; Whitchurch et al., 2004). These four mutants indicate that *P. aeruginosa* uses the Pil-Chp system for a pilus-mediated chemotaxis response to *S. aureus*. Yet how these chemotaxis-deficient mutants control *P. aeruginosa* attraction towards *S. aureus* is not precisely defined by macroscopic directional motility assays.

**Figure 1:**
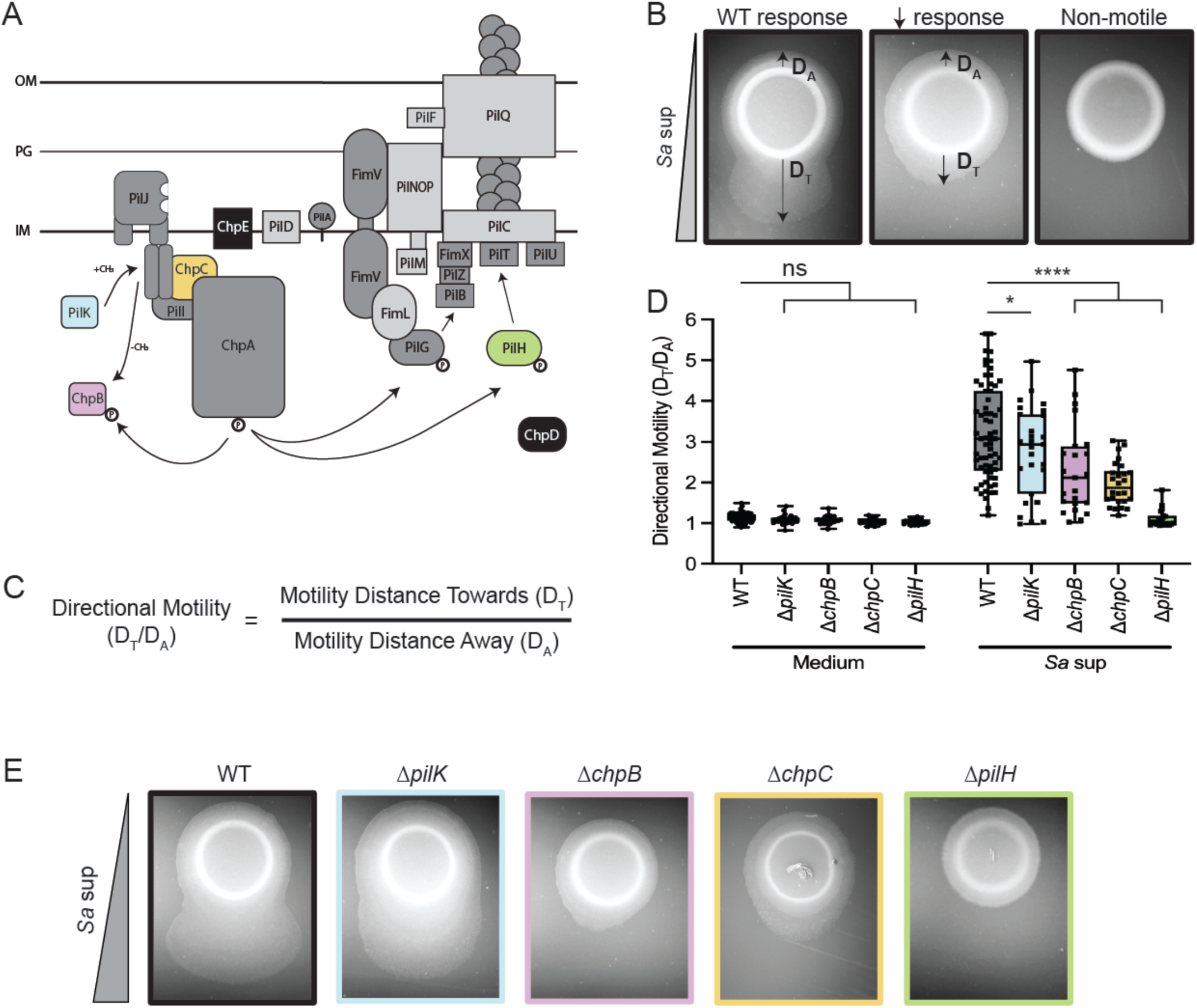
The Pil-Chp system controls *P. aeruginosa* attraction to *S. aureus* peptides. (A) Schematic of the putative type IV pilus (TFP) Pil-Chp chemosensory pathway. When the colored proteins are deleted, cells retain TFP motility, but show reduced pilus-mediated response to the gradient of *S. aureus* secreted factors. Mutants for the proteins highlighted in black retain TFP motility and wildtype levels of response up a gradient of *S. aureus* secreted factors. When the proteins highlighted in dark gray are deleted, the cells are non-motile. Proteins highlighted in light gray were not tested here, but have been previously reported to lack TFP motility. Representative images for each response type are shown in (B). (C) Response measured by calculating the ratio of directional motility up the gradient of *S. aureus* secreted factors. The equation for calculating directional motility is shown. (D) Directional motility of *pil-chp* mutant candidates that retain TFP motility, but show reduced response to *S. aureus* supernatant and representative directional motility images for the wildtype and each mutant shown in (E). Directional motility for at least three biological replicates, each containing a minimum of four technical replicates are shown and statistical significance determined with a two-way ANOVA followed by Dunnett’s multiple comparisons test. *\*\*\*\** indicates p< 0.0001; * indicates p<0.05; *ns* indicates no statistically significant difference in directional motility compared to wildtype *P. aeruginosa*.

## Results: Methyl modification proteins for chemotaxis adaption are necessary for full directional TFP-mediated motility towards *S. aureus*

Since both predicted adaption proteins for chemoreceptor methylation modification are necessary for full response to *S. aureus* peptides at the community level, we next investigated the behavior of each mutant at the single-cell level to uncover how each mutant is unable to correctly bias the direction of movement. PilK is predicted to methylate, while ChpB is predicted to demethylate, PilJ (Figures 1A, 2A). For other MCPs, the addition and removal of methyl groups to a chemoreceptor, facilitates intracellular signal transmission to the downstream kinase to shift activity to an ‘ON’ or ‘OFF’ state, while also inducing conformational changes to the ligand binding region of the MCP that alters its sensitivity to signals as the bacterial cell moves up or down a gradient (Parkinson et al., 2015). Higher levels of MCP methylation, facilitated by PilK, are thus expected to shift kinase activity ON and subsequently increase Pil-Chp control of pilus extension and retraction, while hydrolysis of methylated sites by ChpB is expected to shift kinase activity towards an OFF state, with a reduction in controlled pilus dynamics (Figure 2A).

**Figure 2.**
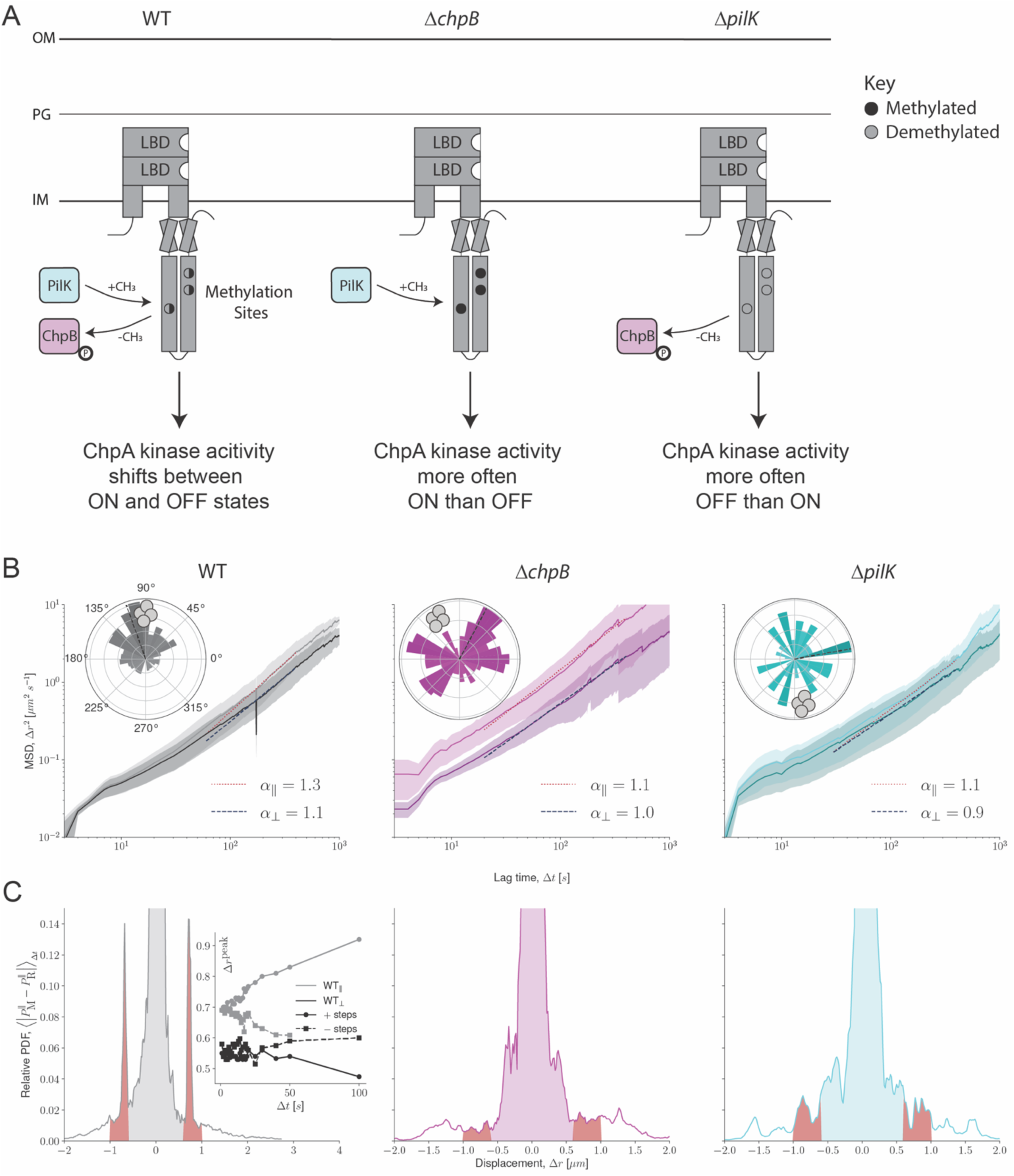
Methyl modification proteins for chemotaxis adaption are necessary for full directional TFP-mediated motility towards *S. aureus.* (A) Schematic of *P. aeruginosa* MCP protein PilJ and methyl modification proteins PilK and ChpB. The predicted methylation sites with cytoplasmic domain of PilJ are represented by the black and gray circles. In wildtype, PilJ is predicted to undergo methylation and demethylation; thus, ChpA kinase activity switches between ON and OFF states (left). In the absence of *chpB*, PilJ is expected to have high methylation (filled, black circles) and therefore ChpA kinase activity shifted ON (middle). In the absence of *pilK*, PilJ is expected to have low methylation (empty, gray circles) and therefore ChpA kinase activity shifted OFF (right). (B) Mean-squared displacements (MSD) for the parallel (||) and perpendicular (⊥) directions of wildtype, *ΔchpB,* and *ΔpilK*. The anomalous diffusion exponent (*α*) for each MSD is shown. Insets show the rose graphs of the principal angle of motility for each cell trajectory relative to starting position for *P. aeruginosa* wildtype, *ΔchpB*, and *ΔpilK* in coculture with *S. aureus.* Position of *S. aureus* relative to the center of the *P. aeruginosa* microcolony is represented by the gray cocci on the perimeter. Trajectory angles are shown by colored vectors with the average angle of all trajectories represented by the dotted black line. Larger vectors indicate more cells for the given principal angle. One rose graph representative of at least three replicates is shown. (C) Relative distributions of parallel component step displacements (*Δr*) of wildtype, *ΔchpB*, and *ΔpilK* cells. Relative distributions are the absolute difference between probability density functions (PDF) of movers and resters, averaged over lag times *Δt ≤ 20* seconds. The red regions highlight the non-zero sharp-shouldered peak step size between *0.6 µm ≤ |Δr| ≤ 1.0 µm*. The inset shows the location of the peak step size for wildtype movers over increasing lag times for forward and backwards steps in both parallel and perpendicular directions.

Based on the information above, we predicted that cells lacking PilK would have low methylated PilJ, while those lacking ChpB would have highly methylated PilJ. Thus, both are predicted to be unable to properly transmit intracellular signals to control activity of kinase ChpA and subsequently the response regulators, ultimately resulting in altered control over biased extension and retraction events for motility (Figure 2A). Macroscopically, the *chpB* mutant, which presumably has high methylation and thus a shifted-ON kinase state, exhibits a decrease in directional movement up the *S. aureus* secreted factor gradient which is restored when *chpB* is complemented (Figure 1, Figure 2 – figure supplement 1). To uncover how *ΔchpB* fails to move fully towards *S. aureus* signals, we evaluated single-cell TFP-mediated motility behaviors of *ΔchpB* in the absence and presence of *S. aureus.* Compared to wildtype *P. aeruginosa,* in both monoculture and coculture, the *ΔchpB* mutant exhibits earlier motility away from the growing microcolony, with groups of motile cells migrating outwards in all directions (Videos 1-4). In coculture with *S. aureus,* some cells migrate towards *S. aureus*; however, similar numbers of cells move in the opposite directions, suggesting *ΔchpB* cells are unable to bias movements towards the interspecies signals, unlike wildtype, which shows stronger apparent bias with more cells moving towards *S. aureus* (Video 4). To quantify these behaviors, we determined the direction of motion for each *P. aeruginosa* cell in relation to the position of *S. aureus* cells. Rose graphs of the principal angles of motility for each *P. aeruginosa* cell trajectory, reveal that most wildtype cells bias their collective direction of movement towards *S. aureus*, whereas *ΔchpB* cells are prone to move both towards and away from *S. aureus* (Figure 2B, inset).

To determine the aspects of *P. aeruginosa* movement that contribute to biased motion towards *S. aureus,* the dynamics were quantified by the mean-squared displacement (MSD), a measure of the typical distance traveled over a given time *Δt* (Figure 2B). While MSD measurements commonly assume isotropic motion, we decomposed microscopic cell migration into components parallel (||) and perpendicular (⊥) to each trajectory’s principal direction. While wildtype cells travel comparable distances in the parallel and perpendicular directions, only significantly different at the longest lag times, they have different dynamics in the two directions (Figure 2B, left). Perpendicular to the principal direction, the cells perform an unbiased random walk, which is quantified by fitting the anomalous diffusion exponent (Methods and Materials; Eq 4) and finding *α⊥≈1.1*, which is close to the expected value of 1 for diffusive dynamics. However, in the parallel direction, the anomalous exponent *α||=1.3*. This larger value suggests that cells tend to move with more self-directed, propulsive transport in the parallel direction.

In comparison to wildtype, cells lacking ChpB exhibit markedly different MSD profiles. *ΔchpB* cells have significantly larger displacements in the parallel direction than perpendicular (Figure 2B, middle). However, this increased microscopic motion does not translate into directional motility, since the anomalous diffusion exponent is near unity, *α||=1.1*, which suggests diffusive random walk dynamics. The combination of increased MSD but loss of self-directed, propulsive transport compared to wildtype is consistent with ChpA kinase activity more often in the ON state. Given the predicted chemotactic role for Pil-Chp, one possible explanation for reduced bias towards *S. aureus* is that *ΔchpB* cells have increased rates of cellular reversals. Reversing, or changing the direction of type IV pilus-mediated movements, is thought to require switching of the leading and lagging poles. These cellular reversals also enable bacteria to bias the direction of type IV pilus-mediated movement up an increasing concentration gradient of chemoattractant (Oliveira et al., 2016). However, single-cell tracking analyses under these conditions did not show statistically significantly different reversal dynamics between *ΔchpB* and wildtype. Therefore, if pole switching reversals do not explain how cells can perform TFP-mediated chemotaxis towards interspecies signals, then how are cells able to bias their movements up a gradient?

To understand how directional motility arises from the microscopic dynamics, we considered the displacement distribution function, which gives the probability that a cell moves a given displacement over a given lag time (Figure 2 - figure supplement 3). The probability density functions of displacements made by *P. aeruginosa* cells are dominated by a narrow peak of submicron jiggling motion. While imaging introduces narrow Gaussian noise (Figure 2 - figure supplement 4A), the submicron peak is observed to be wider and exponentially distributed (Figure 2 - figure supplement 4B). Neither jiggling, nor rare-but-large steps, show substantial differences parallel or perpendicular to the principal direction of motion for wildtype cells. The dynamics leading to directional motility in wildtype cells become apparent once the trajectories are divided into subpopulations of “movers” and “resters” (see Methods and Materials). While rester-designated cells exhibit displacement distributions with only jiggling, movers possess a pair of non-zero sharp-shoulder peaked step sizes in the direction parallel to the principal angle (Figure 2C, red regions). These peaks represent a well-defined step size of 0.69 ± 0.01 µm at the shortest lag times. For wildtype cells, these sharp peaks are found at step sizes of nearly one micron for all lag times and exist in both the parallel and perpendicular directions but are entirely absent in resters. The peak step size is larger in the parallel direction than the perpendicular direction (Figure 2 - figure supplement 3, top row), explaining the anisotropic motion and single-cell directional motility, but not bias towards *S. aureus*. The bias occurs because the symmetry between the forward and backward peaks breaks with increasing lag time, with the forward (positive, parallel) peak shifting to slightly larger step sizes and the backward (negative, antiparallel) peak shifting to slightly smaller values and eventually vanishing at the largest lag times (Figure 2C, left, inset). The probability distributions at different time lags reveal the microscopic basis of the directed motility of wildtype *P. aeruginosa* towards *S. aureus* and the difference between these peak step sizes explains the superdiffusive anomalous exponent. Comparison of wildtype and *ΔchpB* step sizes show that both strains exhibit the dominant submicron jiggling peak with no selected direction much of the time (Figure 2C). However, the peak step size can no longer be distinguished in *ΔchpB* (Figure 2C, middle, red region). This suggests an apparent loss of directional TFP-mediated dynamics and is consistent with the reduced macroscopic directional motility observed in *ΔchpB* (Figure 1D, E).

Given that reduced pilus-mediated chemotaxis of *ΔchpB* can be explained by a loss of the peak step size in the parallel direction, we next live-imaged *ΔpilK* and tracked cells to evaluate whether *ΔpilK* exhibits similar reduction in bias, thus explaining the modest macroscopic chemotaxis deficiency. Observations of *ΔpilK* in monoculture and coculture with *S. aureus* show similar, but bimodal phenotypes. While 50% of *ΔpilK* microcolonies imaged show hypermotile single-cells or small packs of cells moving outwards in all directions from the growing microcolony, regardless of where *S. aureus* is in coculture, the other 50% of imaged *ΔpilK* microcolonies do not show any motility (Figure 2 – figure supplement 2, Videos 5-8). When motile, *ΔpilK* cells disperse from the microcolony later than *ΔchpB* but earlier than wildtype (Video 6). Compared to *ΔchpB* cells that tend to move as clusters of cells in tendril-like patterns away from the microcolony, *ΔpilK* cells more frequently move in smaller groups or strictly as single cells and spread outwards radially (Videos 6 and 8). Regardless, complementation of the *ΔpilK* mutant restores chemotaxis response to wildtype levels (Figure 2 – figure supplement 1). The tracking analyses show that *ΔpilK* MSD is comparable to wildtype but the parallel anomalous exponent *α||=1.1* is closer to the diffusive random walk value of *ΔchpB* (Figure 2B, right). Similarly, like *ΔchpB*, *ΔpilK* has largely lost the peak step size (Figure 2C, right). Compared to *ΔchpB*, the remnant of the peak step size may be discerned (Figure 2C, right, red region); however, they are indefinite and the larger step size of 1.57 ± 0.02 µm at the shortest lag times may also be present (Figure 2 - figure supplement 3, middle row).

Collectively, the bimodal nature of *ΔpilK* motility and phenotype of *ΔchpB* suggest that *P. aeruginosa* cells can lose control of directional pilus-mediated motility dynamics in two ways. Cells with too much ChpA kinase activity, such as *ΔchpB*, move far without precise control over direction and resulting in persistent movement in the initial direction of movement, resulting in larger displacement than wildtype. In contrast, cells with loss of PilJ-mediated ChpA kinase activity, like *ΔpilK,* are unable to control pilus dynamics and thus are more susceptible to environmental conditions, which leads to either the inability to move or uncontrollable, short movements away from the microcolony. This is further exemplified for *ΔpilK* when multiple microcolonies from four separate cultures were simultaneously imaged and retain the bimodal distribution of motility (Figure 2 – figure supplement 2). Whether persisting longer or spreading, neither *ΔchpB* nor *ΔpilK* can chemotax towards interspecies signals, supporting the model that proper methylation and demethylation of chemoreceptor PilJ is an essential component of pilus-mediated chemotaxis.

We previously reported that PilJ was not necessary for *P. aeruginosa* to bias movement towards *S. aureus* (Limoli et al., 2019). However, these data reveal a role for two enzymes predicted to modify PilJ; thus, we revisited the necessity of PilJ in interspecies signaling here. One hypothesis for the prior observations was that motile *ΔpilJ* cells were using flagella-mediated motility, which obscured the pilus-mediated defect. To test this hypothesis, we generated a mutant lacking both *pilJ* and *flgK*, the flagellar hook; therefore, this mutant was not able to use flagella-mediated motility (O’Toole & Kolter, 1998). Live-imaging shows that *ΔpilJ ΔflgK* cells are non-motile and thus fully non-responsive to *S. aureus*, suggesting that previously it was flagella-mediated motility interfering with evaluation of pilus-mediated response to *S. aureus* (Figure 2 – video 1). Additionally, as shown prior with a *ΔpilA ΔflgK* mutant, which similarly lacked functional pili and flagella, *ΔpilJ ΔflgK* cells are unable to remain in a clustered microcolony at later time points (Figure 2 – video 1) (Limoli et al., 2019). To test that *ΔchpB* and *ΔpilK* phenotypes at the single-cell level were not due to flagella-mediated motility as well, we generated *ΔchpB ΔflgK* and *ΔpilK ΔflgK* mutants and live-imaged each in monoculture. Both of these mutants phenocopy the behaviors of their respective parental single mutant strains (Figure 2 – videos 2-4).

**Video 1.** *P. aeruginosa* wildtype in monoculture. Duration 3.5 hr. 3 hr post-inoculation. Acquisition interval 1 sec. Output interval every 40^th^ frame at 50ms/frame.

**Video 2.** *P. aeruginosa* wildtype in coculture. Duration 1 hr 45 min. 3 hr post-inoculation. Acquisition interval 1 sec. Output interval every 40^th^ frame at 50ms/frame.

**Video 3.** *P. aeruginosa ΔchpB* in monoculture. Duration 2 hr. 3 hr post-inoculation. Acquisition interval 1 sec. Output interval every 40^th^ frame at 50ms/frame.

**Video 4.** *P. aeruginosa ΔchpB* in coculture with *S. aureus* wildtype. Duration 3 hr. 2 hr post-inoculation. Acquisition interval 1 sec. Output interval every 40^th^ frame at 50ms/frame.

**Video 5.** *P. aeruginosa ΔpilK* in monoculture. Growing microcolony cells. Duration 3 hr. 3 hr post-inoculation. Acquisition interval 1 sec. Output interval every 40^th^ frame at 50ms/frame.

**Video 6.** *P. aeruginosa ΔpilK* in monoculture. Hypermotile cells. Duration 3 hr. 3 hr post-inoculation. Acquisition interval 1 sec. Output interval every 40^th^ frame at 50ms/frame.

**Video 7.** *P. aeruginosa ΔpilK* in coculture with *S. aureus* wildtype. Growing microcolony cells. Duration 4 hr. 2 hr post-inoculation. Acquisition interval 1 sec. Output interval every 40^th^ frame at 50ms/frame.

**Video 8.** *P. aeruginosa ΔpilK* in coculture with *S. aureus* wildtype. Hypermotile cells. Duration 4 hr. 2 hr post-inoculation. Acquisition interval 1 sec. Output interval every 40^th^ frame at 50ms/frame.

**Figure 2 – video 1.** *P. aeruginosa ΔpilJ ΔflgK* in coculture with *S. aureus* wildtype. Duration 1 hr 15 min. 3 hr post-inoculation. Acquisition interval 1 sec. Output interval every 40^th^ frame at 50ms/frame.

**Figure 2 – video 2.** *P. aeruginosa ΔchpB ΔflgK* in coculture with *S. aureus* wildtype. Duration 3 hr. 3 hr post-inoculation. Acquisition interval 1 sec. Output interval every 40^th^ frame at 50ms/frame.

**Figure 2 – video 3.** *P. aeruginosa ΔpilK ΔflgK* in coculture with *S. aureus* wildtype. Growing microcolony cells. Duration 2.5 hr. 3 hr post-inoculation. Acquisition interval 2 sec. Output interval every 20^th^ frame at 50ms/frame.

**Figure 2 – video 4.** *P. aeruginosa ΔpilK ΔflgK* in coculture with *S. aureus* wildtype. Hypermotile cells. Duration 2.5 hr. 3 hr post-inoculation. Acquisition interval 2 sec. Output interval every 20^th^ frame at 50ms/frame.

## Results: Methyl modification of PilJ is necessary for TFP-mediated chemotaxis response to *S. aureus*

The necessity of PilK and ChpB for directional twitching motility suggests that, like other bacterial chemoreceptors, methylation levels of PilJ influence the downstream signaling cascade; yet, number and location of the methylation sites in PilJ are unknown. We hypothesized that PilJ contained at least one methylation site for chemoreceptor adaptation. To investigate this hypothesis, we first searched the amino acid sequence of PilJ for the conserved ten amino acid methylation motif. This motif, [A/S/T/G]-[A/S/T/G]-X-X-[**E/Q**]-[**E**/**Q**]-X-X-[A/ S/T/G]-[A/S/T/G], has a pair of glutamine and/or glutamate residues at the center (Alexander & Zhulin, 2007; Ortega et al., 2017; Salah Ud-Din & Roujeinikova, 2017; Terwilliger et al., 1986). In other MCPs, one of these residues is methylated by the methyltransferase and demethylated by the methylesterase. Examination of PilJ reveals two motifs with an exact match to the conserved sequence and one motif that shares nine of the ten conserved amino acids. This suggests PilJ has three potential methylation sites in the predicted cytoplasmic region (Figure 3A, black and pale orange circles; Figure 3 – figure supplement 1). The identified methylation sites are located at residues Q_412/E413_, Q_623/Q624_, and Q_639/E640_.

**Figure 3.**
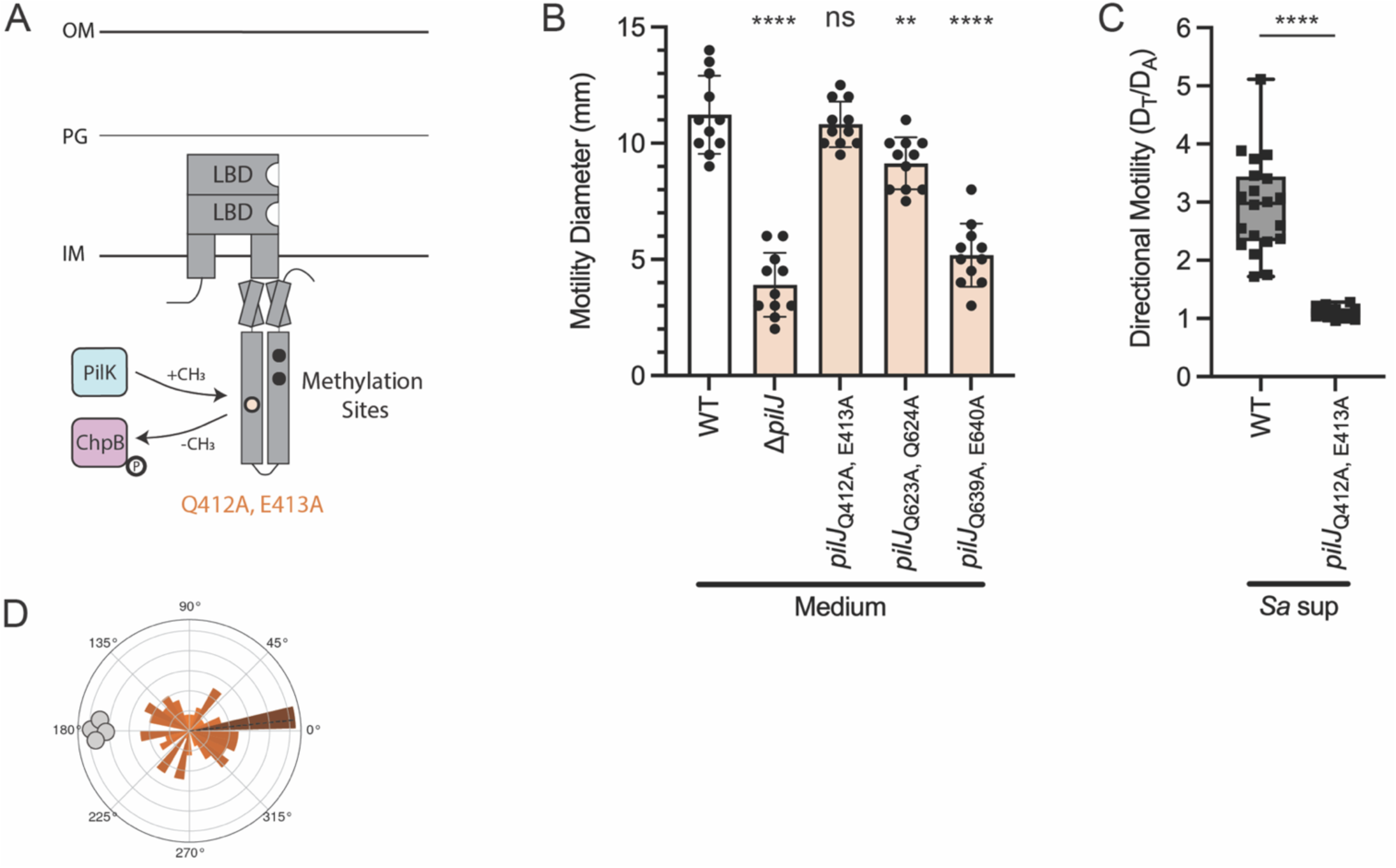
Methyl modification of PilJ is necessary for TFP-mediated chemotaxis response to *S. aureus.* (A) Schematic of PilJ with cytoplasmic methylation sites represented as black circles. The methylation site Q_412, E413_, whose mutant retains wildtype motility and is further studied for chemotaxis response, is highlighted in pale orange. (B) Twitching motility diameters of *P. aeruginosa* wildtype and methylation mutants. Motility diameters were analyzed by a one-way ANOVA followed by Dunnett’s multiple comparisons test. *\*\*\*\** indicates p< 0.0001; ** indicates p<0.01; *ns* indicates no statistically significant difference. (C) Migration towards *S. aureus* secreted factors of *P. aeruginosa* wildtype and methylation mutant *pilJ*Q_412A, E413A_. Directional motility measurements were analyzed with an unpaired t-test. *\*\*\*\** indicates p< 0.0001. Motility diameters and directional motility are shown for at least three biological replicates each containing a minimum of four technical replicates. (D) Rose graph of the principal angle of motility for each cell trajectory relative to starting position for *P. aeruginosa pilJ*Q_412A, E413A_ in coculture with *S. aureus.* Position of *S. aureus* relative to the center of the *pilJ*Q_412A, E413A_ cells is represented by the gray cocci on the perimeter. Trajectory angles are shown by colored vectors with the average angle of all trajectories represented by the dotted black line. Larger vectors indicate more cells for the given principal angle. One rose graph representative of at least three replicates is shown.

To experimentally determine whether the methylation sites were necessary for TFP-mediated chemotaxis, we next generated a mutation in each methylation site by substituting the glutamine/glutamate residue pair to an alanine/alanine pair. Given most other *pilJ* mutants exhibit severely diminished twitching motility, we first compared the ability of each PilJ methylation site mutant to twitch in a standard subsurface motility assay (Turnbull & Whitchurch, 2014). Briefly, *P. aeruginosa* was inoculated at the plastic-agar interface and allowed to move prior to measuring the diameters of motility. Only the *pilJ*Q_412A, E413A_ mutant retains full wildtype levels of motility, while *pilJ*Q_623A, Q624A_ has moderate-but-significant reduction in motility and *pilJ*Q_639A, E640A_ is severely diminished in twitching motility (Figure 3B). As the only methylation site mutant that retains wildtype levels of pilus-mediated motility, the next question is whether *pilJ*Q_412A, E413A_ exhibits reduced attraction towards *S. aureus.* In the macroscopic directional motility assay, *pilJ*Q_412A, E413A_ demonstrates significant loss of migration up the gradient of *S. aureus* supernatant (Figure 3C). The other methylation site mutants also display attenuated chemoattraction to *S. aureus*; yet, it is unclear how much of this reduced response is due to motility defects versus deficiency in pilus-mediated chemotaxis (Figure 3 – figure supplement 2).To confirm that the migration phenotype of *pilJ*Q_412A, E413A_ can be restored with full-length *pilJ,* the mutant and wildtype strains were complemented with a plasmid containing a GFP-tagged copy of wildtype *pilJ* under control of an arabinose-inducible promoter (Figure 3 – figure supplement 3). This allows for visualization of PilJ in the cells, which show the expected bipolar localization (Figure 3 – figure supplement 3A). Additionally, the complemented mutant strain shows levels of directional response similar to wildtype harboring the complementation plasmid (Figure 3 – figure supplement 3B). Due to the GFP tag, some PilJ signaling may be diminished leading to the lower interspecies signal response in the complemented strains.

Next, we live-imaged the *pilJ*Q_412A, E413A_ mutant in monoculture and observe that it moves earlier than wildtype, *ΔpilK,* or *ΔchpB,* with single-cell or small groups of two to three cells traveling together (Video 9). Furthermore, *pilJ*Q_412A, E413A_ does not form a microcolony; rather, the cells begin to twitch and move apart from each other at very low cell density, typically before there are ∼10 cells present. This behavior is recapitulated in the presence of *S. aureus,* with *pilJ*Q_412A, E413A_ additionally showing no bias in movement towards *S. aureus* (Figure 3D, Video 10). The *pilJ*Q_412A,E413A_ mutant cells do become elongated after a few hours compared to wildtype. This slight cell division defect is likely due to the high cAMP levels in *pilJ*Q_412A, E413A_ (see Figure 5). Live-imaging of a *pilJ*Q_412A, E413A_ *ΔflgK* mutant in coculture with *S. aureus* phenocopies the parental *pilJ*Q_412A, E413A_ (Figure 3 – video 1). Together these data identify the methylation sites of PilJ and show that methyl modification of these sites is important for chemotaxis signaling.

**Figure 4.**
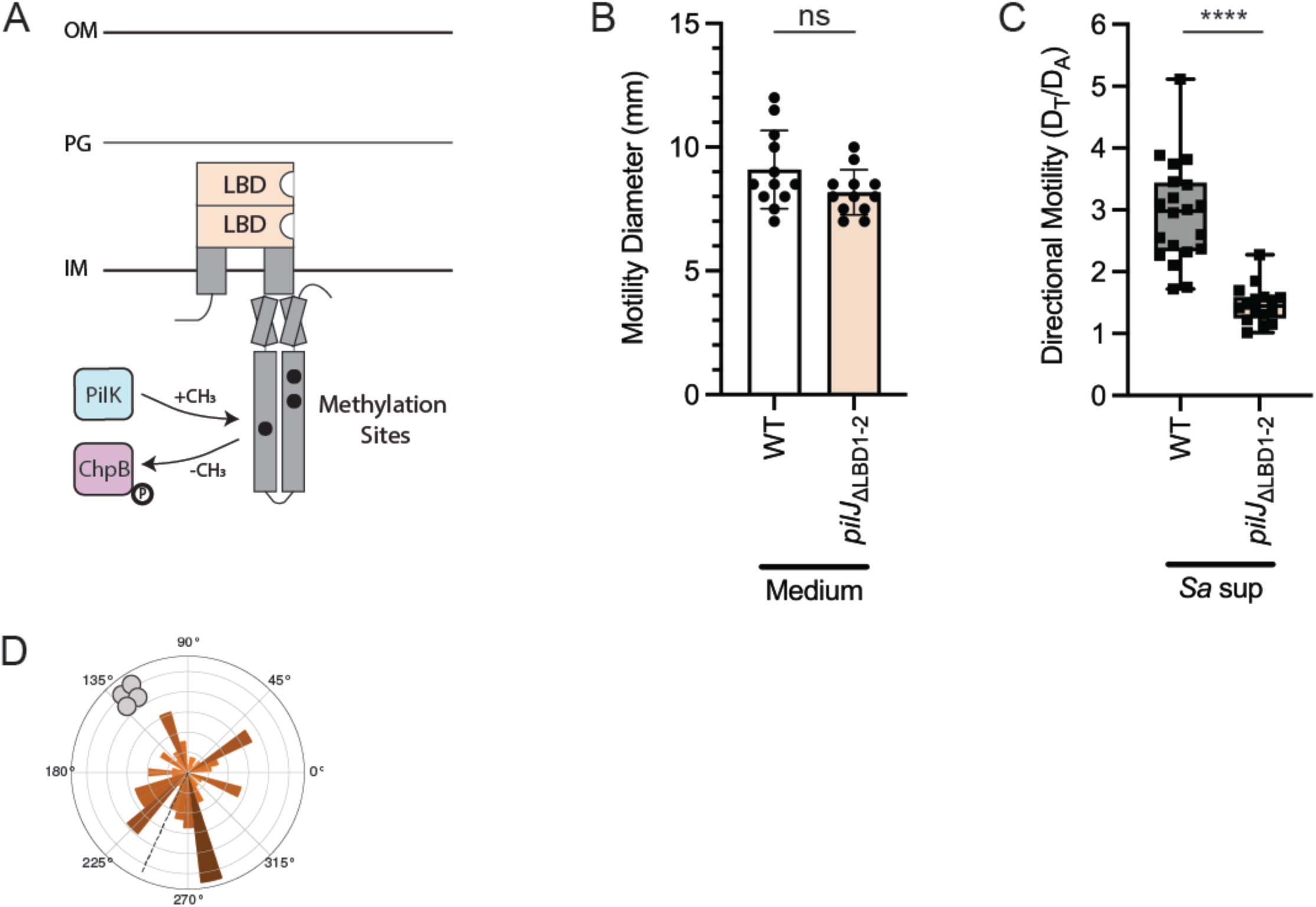
The ligand binding domains of PilJ are required for pilus-mediated chemotaxis but not twitching motility. (A) Schematic of PilJ with periplasmic ligand binding domains (LBDs) highlighted in pale orange. Twitching motility diameters (B) and directional motility towards *S. aureus* secreted factors (C) of *P. aeruginosa* wildtype and *pilJ*_ΔLBD1-2_. Macroscopic motility measurements were analyzed with an unpaired t-test. *\*\*\*\** indicates p< 0.0001; *ns* indicates no statistically significant difference. Motility diameters and directional motility are shown for at least three biological replicates each containing a minimum of three technical replicates. (D) Rose graph of the principal angle of motility for each cell trajectory relative to starting position for *P. aeruginosa pilJ*_ΔLBD1-2_ in coculture with *S. aureus.* Position of *S. aureus* relative to the center of the *pilJ*Q_412A, E413A_ cells is represented by the gray cocci on the perimeter. Trajectory angles are shown by colored vectors with the average angle of all trajectories represented by the dotted black line. Larger vectors indicate more cells for the given principal angle. One rose graph representative of at least three replicates is shown.

**Figure 5.**
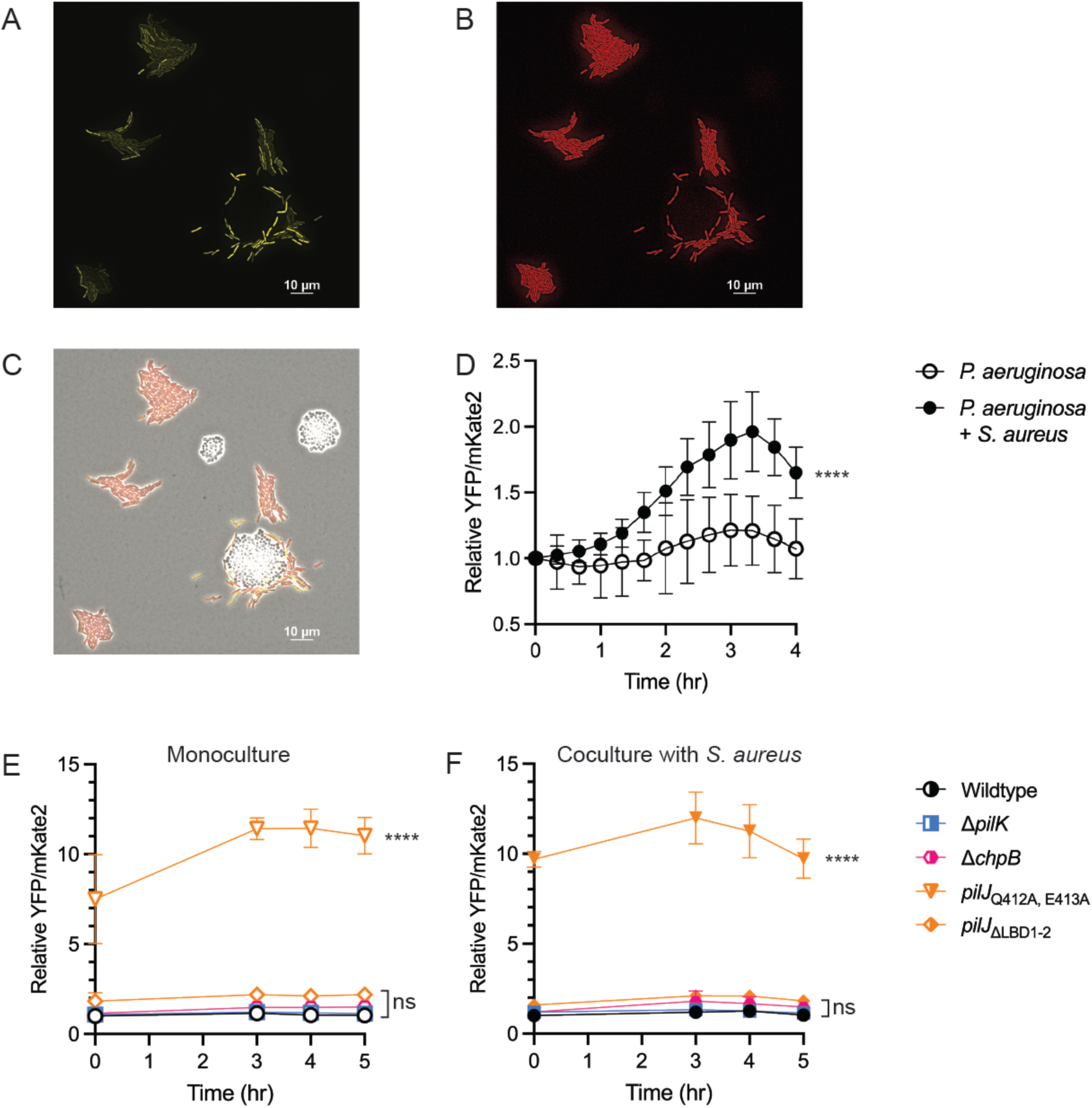
cAMP changes during pilus-mediated chemotaxis independent of TFP activity. (A-D) Intracellular levels of *P. aeruginosa* cAMP measured in monoculture and coculture with *S. aureus* using a *P. aeruginosa* PAO1 strain carrying a reporter with the cAMP-responsive promoter P*xphA* transcriptionally fused to *yfp* and constitutively expressed promoter P*rpoD* fused to *mKate2* for normalization. Representative coculture images at t = 3.5 hr for the YFP (A) TxRed (B), and merged channels (C). cAMP levels of *P. aeruginosa* PA14 *pil-chp* mutants in monoculture (E) and coculture with *S. aureus* (F) using the cAMP reporter. cAMP levels were monitored by dividing YFP by mKate2 fluorescence intensity for each time point and normalizing intensity to wildtype cAMP at t = 0 hr. cAMP levels are shown for at least four microcolonies per condition and were compared using either multiple unpaired t-tests or a one-way ANOVA followed by Dunnett’s multiple comparison’s test. **** indicates p<0.0001; *ns* indicates no statistically significant difference.

**Video 9.** *P. aeruginosa pilJ*Q_412A, E413A_ in monoculture. Duration 2 hr 40 min. 2 hr 20 min post-inoculation. Acquisition interval 1 sec. Output interval every 40^th^ frame at 50ms/frame.

**Video 10.** *P. aeruginosa pilJ*Q_412A, E413A_ in coculture with *S. aureus* wildtype. Duration 3 hr. 2 hr post-inoculation. Acquisition interval 1 sec. Output interval every 40^th^ frame at 50ms/frame.

**Figure 3 – video 1.** *P. aeruginosa pilJ*Q_412A, E413A_ *ΔflgK* in coculture with *S. aureus* wildtype. Duration 3.5 hr. 1 hr post-inoculation. Acquisition interval 1 sec. Output interval every 40^th^ frame at 50ms/frame.

## Results: The ligand binding domains of PilJ are required for pilus-mediated chemotaxis but not twitching motility

Given methylation adaptation is necessary to bias movement towards *S. aureus* but not twitching, we next asked if the LBDs of *P. aeruginosa* PilJ are also required for directing motility. We generated a mutant lacking the periplasmic portion containing both predicted PilJ LBDs (residues 39-303), based on the predictions by Martín-Mora et al. and confirmed by AlphaFold, yet kept the transmembrane domains and entire cytoplasmic region intact, which we now refer to as *pilJ*_ΔLBD1-2_ (Figure 4A, LBDs in pale orange) (Martin-Mora et al., 2019).

We first tested whether *pilJ*_ΔLBD1-2_ retained any twitching motility using the subsurface twitching assay, as described above, and found that loss of the LBDs does not reduce the ability to twitch (Figure 4B). Since *pilJ*_ΔLBD1-2_ is able to twitch to wildtype levels, we next tested the extent that it could chemotax up a gradient of *S. aureus* secreted factors. Despite being able to twitch, *pilJ*_ΔLBD1-2_ loses nearly all ability to move directionally towards *S. aureus* secreted factors (Figure 4C). When complemented with the same *pilJ-gfp* plasmid described above, *pilJ*_ΔLBD1-2_ exhibits bipolar localization of PilJ; however, only partial restoration of interspecies signal response is observed, despite statistical similarity to wildtype with the complement plasmid (Figure 4 – figure supplement 1). Chemoreceptors are typically grouped in arrays of trimer-of-dimer units with each unit signaling downstream to a kinase (Parkinson et al., 2015). Therefore, despite proper localization, the combination of mutant and GFP-tagged wildtype copies of PilJ for this particular strain may lead to trimers-of-dimers with inefficient signaling for complete directional response (i.e., the *pilJ*_ΔLBD1-2_ allele is partially dominant).

Following examination of community level movement and response, we performed live-imaging of *pilJ*_ΔLBD1-2_ in monoculture to determine how loss of the LBDs impacted single-cell movements. The *pilJ*_ΔLBD1-2_ mutant displays increased and earlier motility, with cells lacking microcolony formation seen by the wildtype (Video 11). Unlike the methylation PilJ mutant, *pilJ*_ΔLBD1-2_ tends to move as groups of cells, with trajectories curving in wide loops (Video 11). These behaviors are echoed in coculture with *S. aureus,* with *P. aeruginosa pilJ*_ΔLBD1-2_ appearing to show no bias towards *S. aureus* microcolonies (Figure 4D, Video 12). This behavior is also phenocopied by a *pilJ*_ΔLBD1-2_ *ΔflgK* mutant (Figure 4 – video 1). Collectively, these data establish a role for the LBDs of PilJ in control of response to interspecies signals, while also showing they are not essential for general pilus-mediated motility.

**Video 11.** *P. aeruginosa pilJ*_ΔLBD1-2_ in monoculture. Duration 2 hr 40 min. 2 hr 20 min post-inoculation. Acquisition interval 1 sec. Output interval every 40^th^ frame at 50ms/frame.

**Video 12.** *P. aeruginosa pilJ*_ΔLBD1-2_ in coculture with *S. aureus* wildtype. Duration 2 hr. 3 hr post-inoculation. Acquisition interval 1 sec. Output interval every 40^th^ frame at 50ms/frame.

**Figure 4 – video 1.** *P. aeruginosa pilJ*_ΔLBD1-2_ *ΔflgK* in coculture with *S. aureus* wildtype. Duration 3 hr. 2 hr post-inoculation. Acquisition interval 1 sec. Output interval every 40^th^ frame at 50ms/frame.

## Results: cAMP changes during pilus-mediated chemotaxis independent of TFP activity

Intracellular levels of the second messenger cAMP and Pil-Chp activity are connected. As signal transduction through Pil-Chp increases, the response regulator PilG directly activates the adenylate cyclase, CyaB, necessary for cAMP synthesis (Fulcher et al., 2010). In turn, cAMP indirectly increases transcriptional expression of *pil-chp* generating a positive feedback loop between the two systems (Fulcher et al., 2010; Wolfgang et al., 2003).

Due to the role of Pil-Chp in perception and reaction to *S. aureus,* we asked whether *P. aeruginosa* increased cAMP levels during interactions with *S. aureus*. To answer this question, we live-imaged a previously established cAMP reporter strain *P. aeruginosa* wildtype PAO1 carrying a cAMP-responsive promoter, P*xphA*, fused to *yfp* and a constitutively expressed promoter P*rpoD*, fused to *mKate2* (Persat et al., 2015). Kinetic cAMP levels were measured in individual cells in monoculture or coculture with *S. aureus*, with YFP normalized to mKate2 fluorescence. cAMP is known to be heterogenous amongst cells in a population. In wildtype *P. aeruginosa* monoculture, this heterogeneity is observed, but with minor changes in cAMP levels over time. However, in coculture, cAMP increases, particularly in cells that move towards and surround *S. aureus* (Figure 5A-D). This suggests that response to *S. aureus* is associated with increases in cAMP. Furthermore, when community level response of *P. aeruginosa* lacking either CyaB or the cAMP phosphodiesterase, CpdA, was examined, *P. aeruginosa* cannot fully move up a gradient of *S. aureus* supernatant, despite retaining twitching motility (Figure 5 – figure supplement 1). These data suggest that *P. aeruginosa* controls cAMP levels for proper TFP-mediated response to interspecies signals.

Next, we asked how cAMP levels compared between response-deficient mutants and the parental *P. aeruginosa* (PA14). While reporter levels in PA14 are lower in comparison to PAO1, a similar trend is observed. Each mutant shows at least some increase in cAMP relative to wildtype cAMP levels in monoculture or coculture, though not significant for *ΔpilK* and *ΔchpB*. The *pilJ*_ΔLBD1-2_ mutant exhibits an approximately two-fold increase in cAMP over time compared to wildtype; whereas, *pilJ*_ΔQ412A, E413A_ exhibits a ten-fold increase, independent of the presence of *S. aureus* (Figure 5E, F). These data suggest that cAMP levels are determined by chemoreceptor-mediated signaling, although the degree to which cAMP increases does not always directly correlate with the extent of increased motility.

## Discussion

While the Pil-Chp system has been studied for its roles in twitching motility and cAMP regulation, less work has explored the additional roles of the system in TFP-mediated chemotaxis. Furthermore, most studies have primarily investigated PilJ through a non-motile complete-deletion mutant, which allowed little understanding of the domains of PilJ that determine chemoreceptor activity. Here, we utilized domain-specific mutations of PilJ to evaluate single-cell and community level behaviors that define the importance of PilJ to sense and relay interspecies signals to pilus response regulators. These observations show *P. aeruginosa* is able to move towards interspecies signals through a novel TFP-mediated chemotaxis mechanism and that PilJ does have the necessary components to serve as a MCP for signal sensation, transmission, and adaptation (Figure 6). To our knowledge, this is the first study to generate mutants in the LBDs and methylation sites of PilJ to define their contribution to chemotactic regulation.

**Figure 6.**
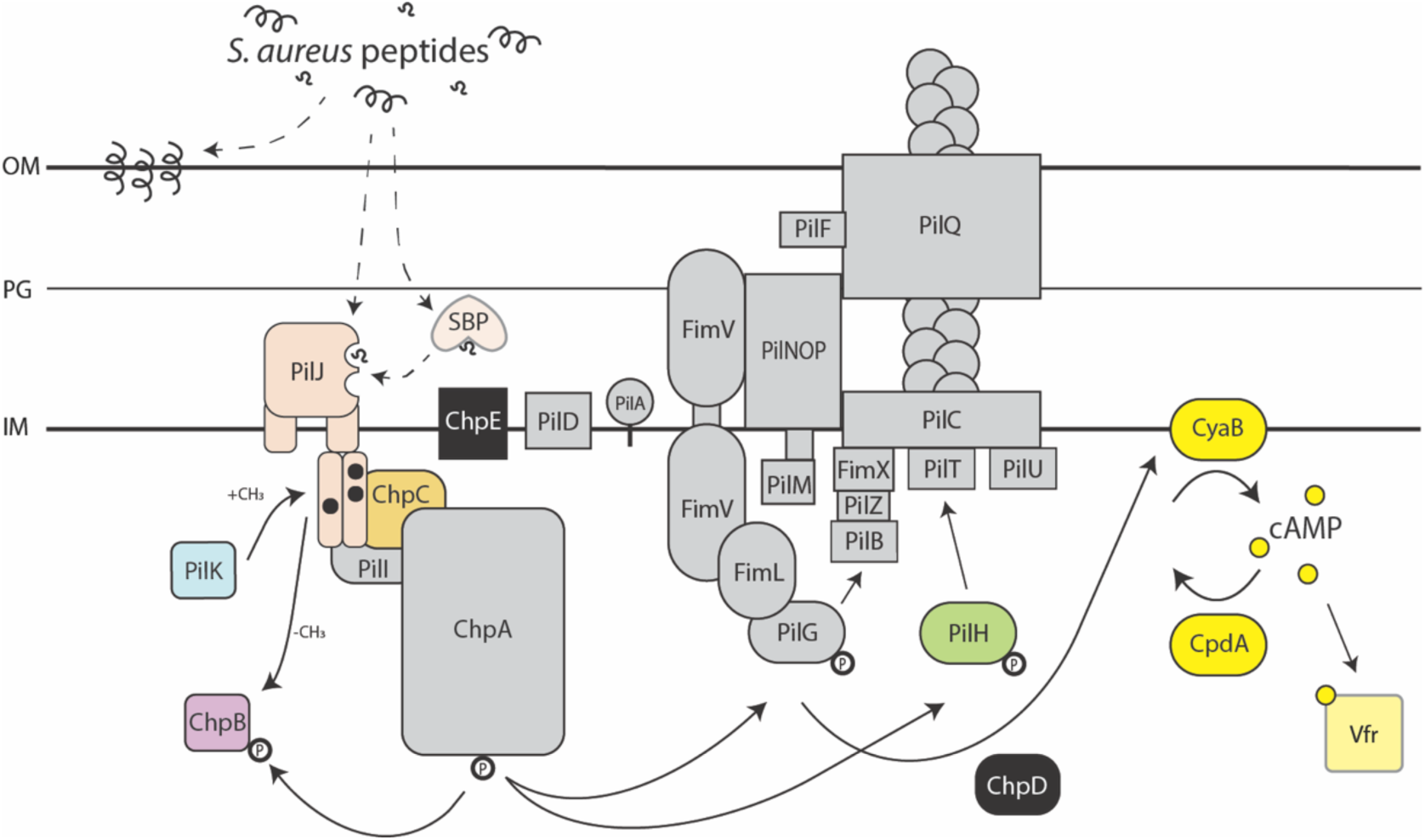
Model of TFP-mediated interspecies chemotaxis. *S. aureus* peptide signals (black) are sensed by *P. aeruginosa* Pil-Chp chemoreceptor PilJ (pale orange) ligand binding domains and transmitted to histidine kinase ChpA (gray), likely through the linker protein ChpC (gold). The kinase state modulates the activity of response regulators PilG (gray) and PilH (green), which coordinate TFP extension and retraction events that bias movements towards the interspecies signals. Adaption of PilJ through modification of the methylation sites by enzymes, PilK (blue) and ChpB (purple), is also required for proper chemotactic response. Chemotaxis through the Pil-Chp system also leads to increases in cAMP (yellow) and depends on careful regulation of these cAMP levels by enzymes CyaB and CpdA (yellow). cAMP activates Vfr for upregulation of virulence factors, including the Pil-Chp system. The chemoattractant peptide signals sensed by PilJ may be full-length *S. aureus* PSMs (black helices) or oligopeptide cleavage products (black squiggles). Additionally, these peptides, whether whole or cleaved, could directly bind PilJ or indirectly activate PilJ via an interaction with a periplasmic solute binding protein (SBP). Alternatively, *S. aureus* peptides may embed into the *P. aeruginosa* membrane, causing transient membrane stress and activating Pil-Chp. The solid arrows indicate previously described interactions and the dashed arrows indicate hypothetical pathways that will be tested in future studies. OM = outer membrane, PG = peptidoglycan, IM = inner membrane.

While flagella-mediated chemotaxis of swimming bacteria has been extensively characterized, there is comparatively little insight into bacterial chemotaxis on a surface. Despite overall sequence similarity of PilJ in the cytoplasmic region to other *P. aeruginosa* MCPs, the low similarity of the periplasmic region containing the LBDs has led to several questions about the function of this putative pilus MCP. First, what ligands does PilJ bind? Martín-Mora et al. previously investigated the only other *P. aeruginosa* MCP containing a PilJ LBD, McpN, which was shown to bind nitrate (Martin-Mora et al., 2019). Evaluation of the McpN and PilJ ligand binding pocket motifs showed little conservation between the two MCPs and PilJ lacked nitrate binding ability (Martin-Mora et al., 2019). If PilJ does not bind nitrate, then what signals can it bind? Persat et al. showed that PilJ can interact with the pilus monomer PilA in the periplasmic regions of each protein and proposed this interaction regulates mechanosensing (Persat et al., 2015), although recent data argues against this model (Kuchma & O’Toole, 2022). *P. aeruginosa* has a second non-conventional chemosensory system for surface sensing, called Wsp. Recently, it was shown that the Wsp system is more broadly a membrane stress detection system and surfaces are just one of many membrane stressors that the receptor WspA detects (O’Neal et al., 2022). In eukaryotic cells, phenol soluble modulins (PSMs) are known to form a membrane-perturbing pore; thus, PilJ may also perceive and transduce interspecies peptide-induced membrane stress (Figure 6) (Verdon et al., 2009).

If PilJ binds a conventional chemoattractant signal, full length PSMs are unlikely to interact directly with the LBD, due to their size and amphipathic secondary structure. Nolan et al. identified that *P. aeruginosa* exhibits increased twitching in the presence of environmental signals, such as tryptone, mucins, or bovine serum albumin (Nolan et al., 2020). This response required *P. aeruginosa* protease activity, presumably to cleave environmental factors into smaller signals (Nolan et al., 2020). While this group did not study the chemotactic nature of these compounds, they do suggest that the increased twitching response is PilJ-mediated. This indicates that PilJ may sense a broad range of environmental signals. It is thus plausible that the *S. aureus* peptides, which are shown to elicit a PilJ-dependent chemotaxis response here, are indeed a chemoattractant signal. Yet, it is still unclear how PSMs may gain access to and bind the periplasmic LBDs of PilJ and thus act as a traditional chemoattractant signal for activation of downstream signaling. Therefore, protease-cleavage of interspecies peptides may be necessary to fragment PSMs into signal-sized peptides for either direct or indirect PilJ binding, in conjunction with earlier observations that *P. aeruginosa* chemotaxes towards di- or tripeptides rather than larger oligopeptides (Kelly-Wintenberg & Montie, 1994). Another possibility is that the peptides bind PilJ indirectly, perhaps through a solute binding protein, as previously reported for chemoattractant inorganic indirectly binding chemoreceptor CtpL through the mediating solute binding protein PtsS (Matilla et al., 2021; Rico-Jiménez et al., 2013). Investigation of these hypotheses, including a role for membrane stress, is currently underway (Figure 6).

Jansari et al. previously reported that their *pilJ* mutant retained some twitching motility, but had diminished response to phosphatidylethanolamine (Jansari et al., 2016). In our current study, chemoattraction to phosphatidylethanolamine by wildtype *P. aeruginosa* was not observed and therefore chemotaxis towards phosphatidylethanolamine by the *pilJ* mutants generated here could not be determined. Jansari et al. generated a *pilJ* mutant lacking residues 74-273 (Jansari et al., 2016). Examination of a similar mutant lacking residues 80-273 in the present investigation shows that *P. aeruginosa pilJ*Δ80-273 similarly has reduced twitching; yet, this defect unfortunately yields insufficient motility to measure reduced attraction to *S. aureus* and, thus, could not be evaluated here (Figure 4 – figure supplement 2). However, the mutant lacking both PilJ domains (residues 39-303; *pilJ*_ΔLBD1-2_) retains near wildtype levels of motility and allows for visualization of response deficiency at both the community and single-cell levels. These observations show that without the PilJ LBDs, *P. aeruginosa* is unable to bias motility towards interspecies signals, which further supports that the periplasmic portion of PilJ is not essential for twitching motility but is important for sensing signals—whether they are chemoattractants or surface signals. This suggests PilJ has evolved to coordinate TFP motility in response to several environmental factors. It remains unknown, however, why *P. aeruginosa* PilJ contains two pilJ LBDs and whether they bind ligands cooperatively or independently, or if each has a designated role for particular chemo- or surface-sensing signals.

Little was previously known about the methylation sites on PilJ, which are typically required for chemotaxis adaptation. While bacterial chemoreceptors have a conserved motif for methylation sites, potential motifs on PilJ had not been identified. This work is the first to identify three likely residue pairs for methyl modification on PilJ, further characterizing the chemoreceptor. In other bacterial chemoreceptors, mutation of each methylation site on a chemoreceptor does not lead to the same phenotype (Astling et al., 2006). Thus, it is reasonable that mutations in each PilJ methylation site differentially influence the conformation of PilJ signaling domains and lead to the three different macroscopic motility behaviors described here (Figure 3B; Figure 3 – figure supplement 2). While live-imaging was only performed on the first methylation site, this was sufficient to show changes in these sites can dramatically alter *P. aeruginosa* chemotaxis (Video 9).

Increases in *P. aeruginosa* cAMP are commonly associated with surface sensing and Pil-Chp activity. It is further established that while PilG activates CyaB for cAMP production, cAMP in turn increases expression of *pil-chp* genes (Fulcher et al., 2010; Wolfgang et al., 2003). However, previous reports focused on Pil-Chp activity in terms of twitching motility; thus, a link between chemotactic Pil-Chp activity and cAMP levels had been ambiguous. While each mutant shown here has varying levels of motility, all are increased in both motility and cAMP levels relative to wildtype; yet, there is not a direct correlation between mutants that have increased motility and the degree to which cAMP is increased. Only *pilJ*Q_412A, E413A_ showed much higher levels of cAMP at nearly 10-fold the amount as wildtype. While the precise role of cAMP signaling in chemotaxis is unclear, studies are currently underway to further interrogate this signaling pathway.

Dissection of Pil-Chp and its role in chemotaxis towards interspecies signals has broadened understanding of a unique bacterial chemosensory system that may be utilized for bacterial communication and survival in complex, polymicrobial environments. During *P. aeruginosa-S. aureus* coinfections, such as those in cystic fibrosis airways, patients often succumb to worse clinical outcomes than their counterparts who are only infected by one organism (Limoli & Hoffman, 2019). Furthermore, once coinfected, patients tend to stay infected by both organisms for several years (Fischer et al., 2021). Such stable, long-term polymicrobial infections may be enhanced by the chemical and physical interactions between species seen here. With this knowledge of how bacteria can sense their respective secreted factors, new therapeutic strategies targeting this system may provide the opportunity to break communication between species and prevent these detrimental interactions thereby eliminating infections and consequently improve patient outcomes.

## Key Resources Table

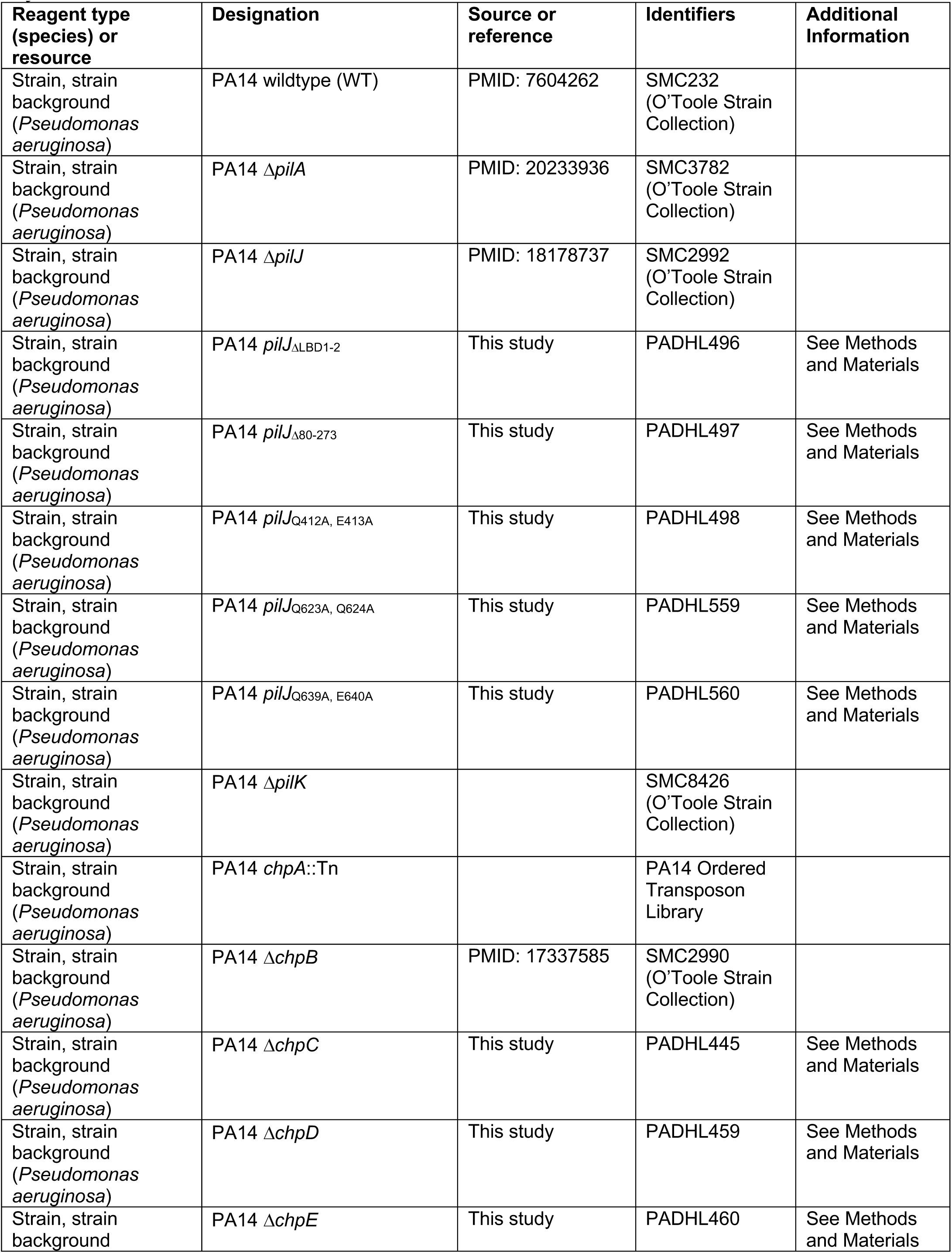
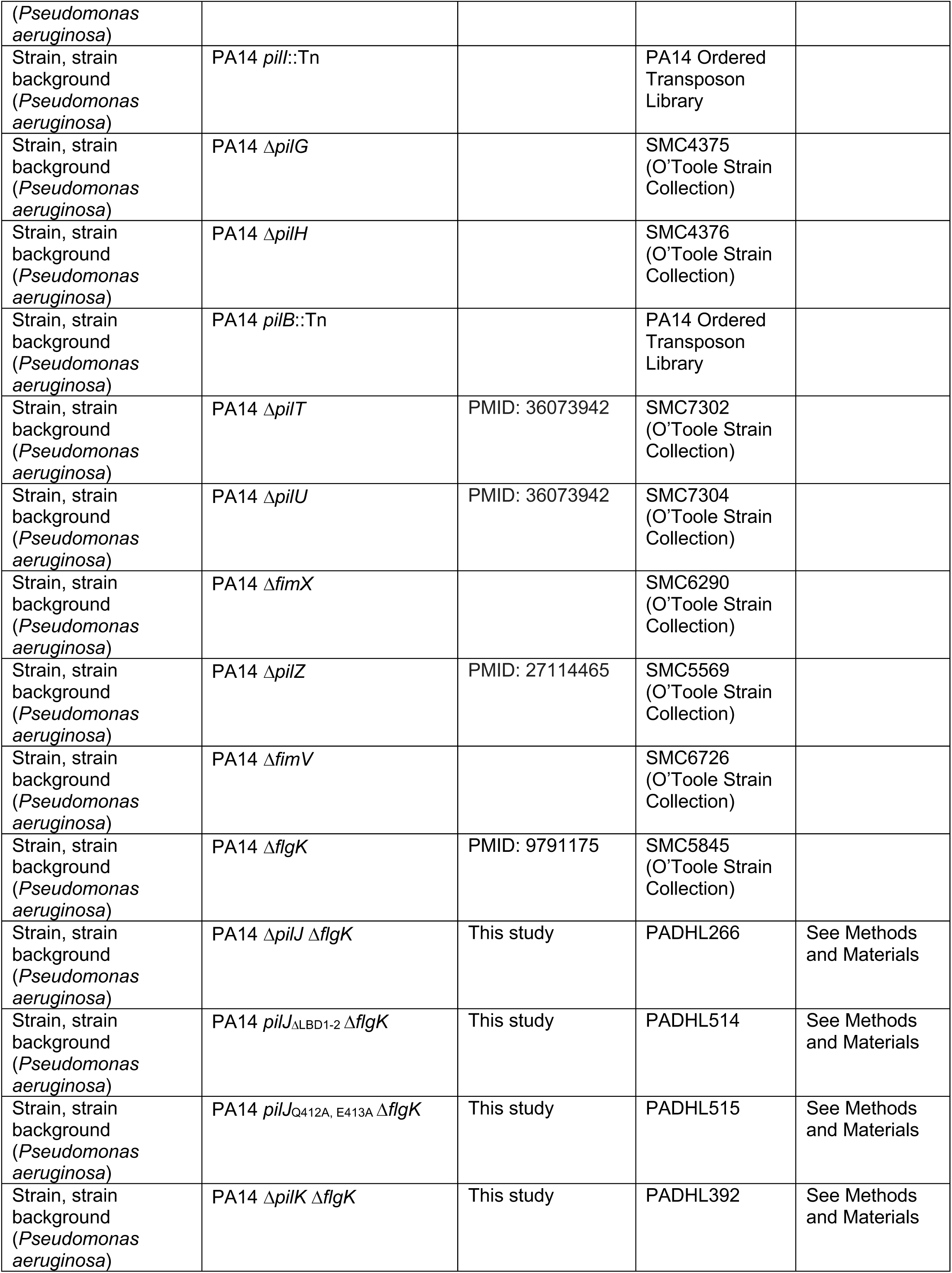
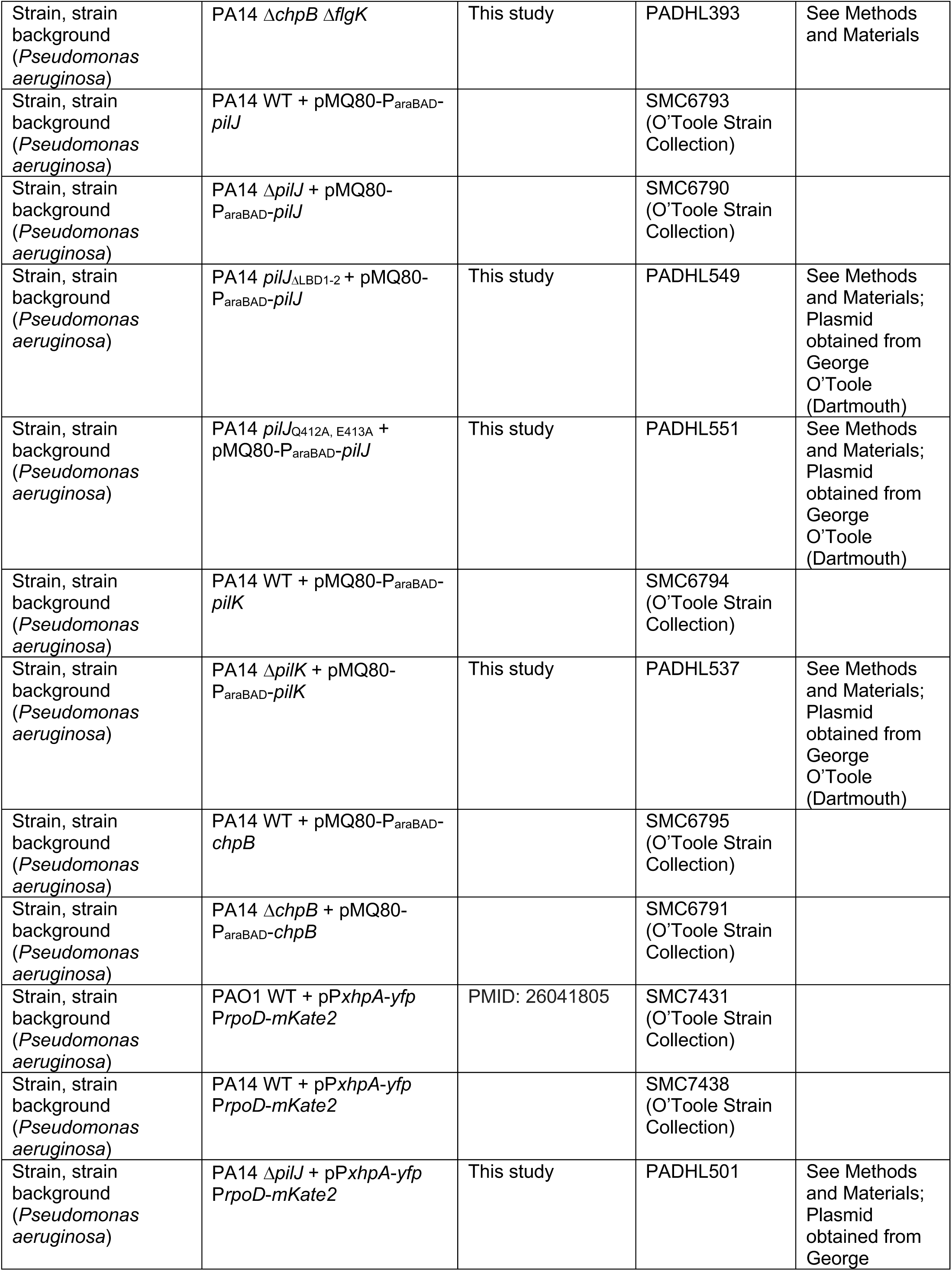
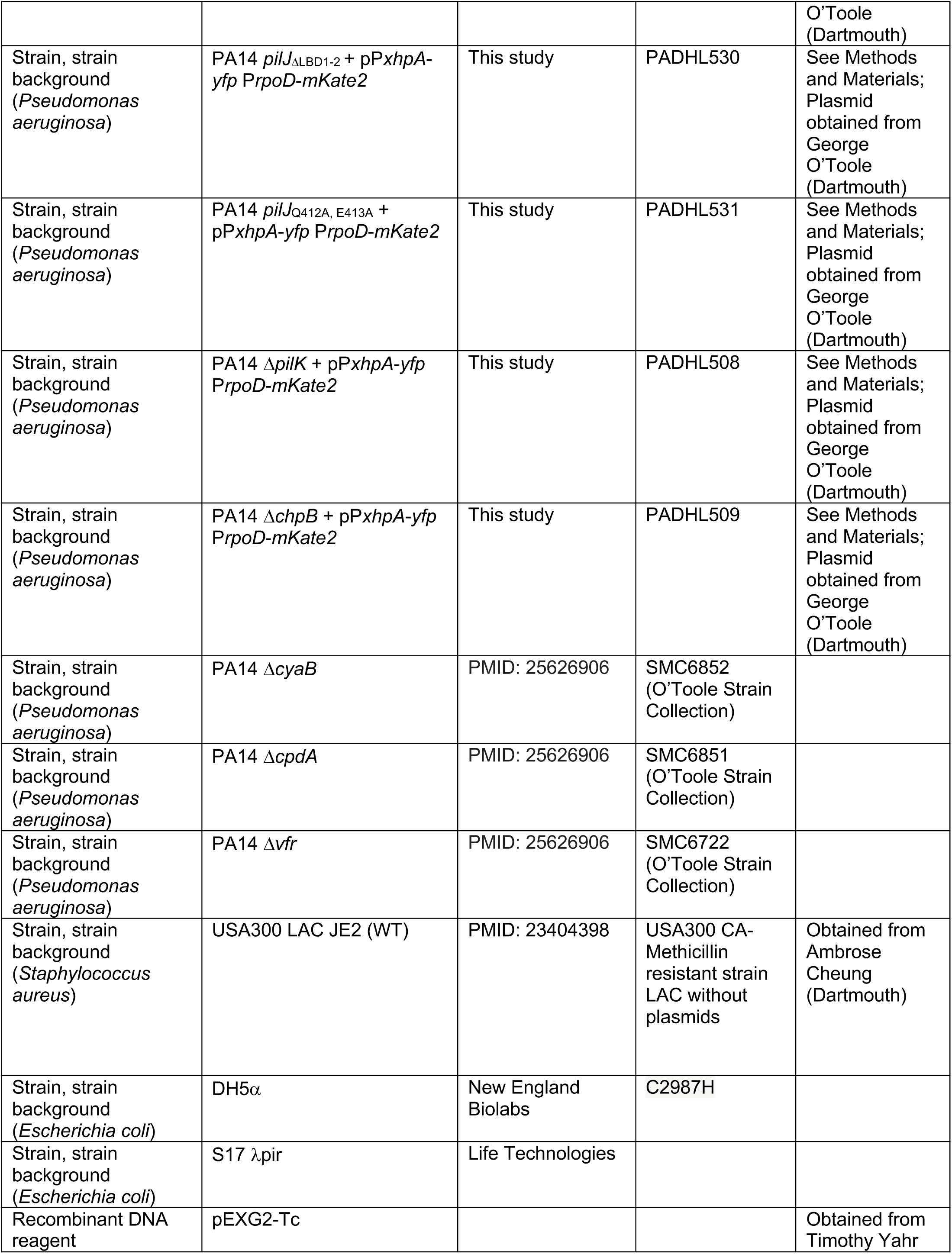
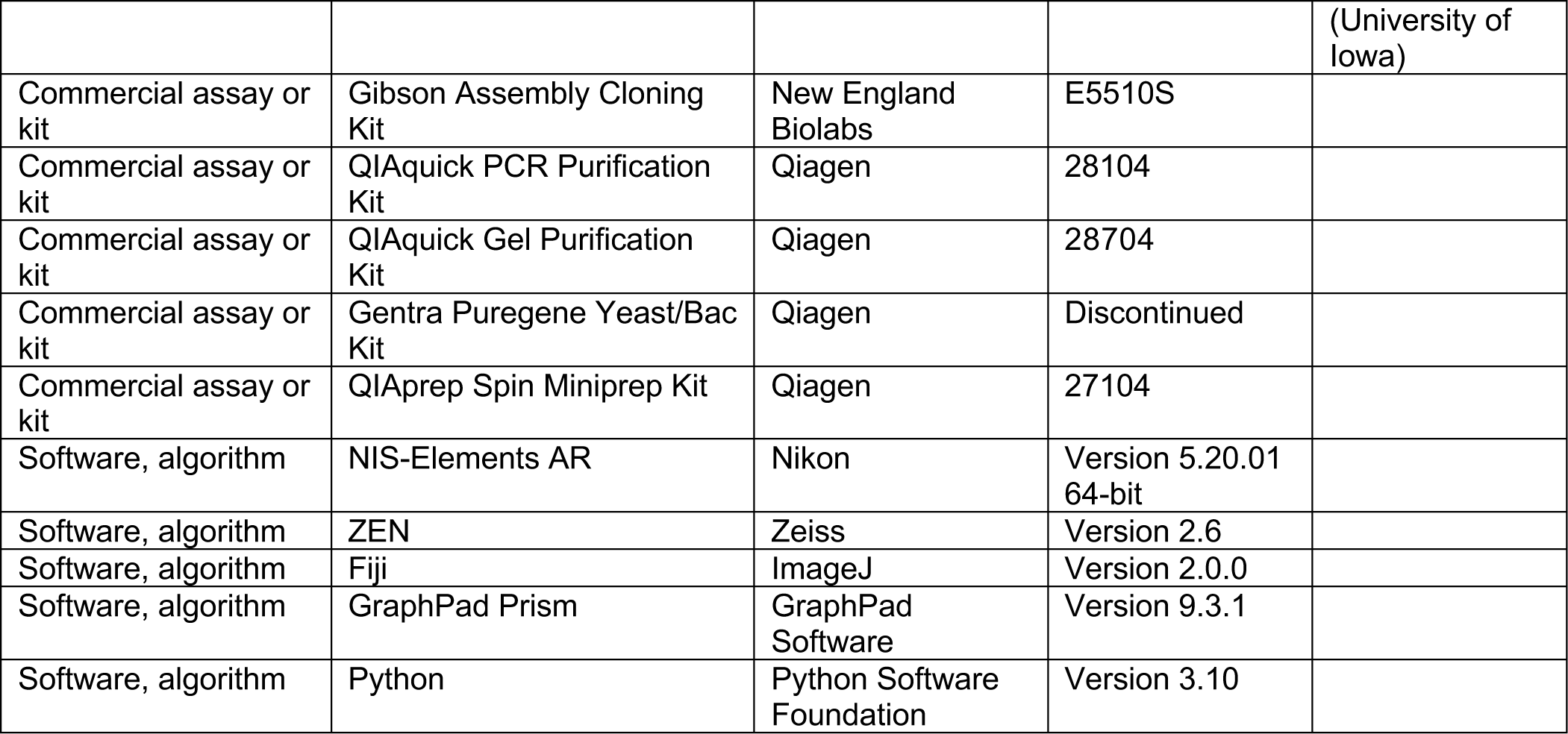

## Methods and Materials

### Bacterial strains and culture conditions

*(i) P. aeruginosa* PA14 or PAO1 and *S. aureus* JE2 strains were cultured in tryptic soy broth (TSB; Becton Dickenson) or M8 minimal media broth supplemented with 0.2% glucose and 1.2% tryptone (M8T) with aeration at 37°C. The following antibiotics were added for *P. aeruginosa* cultures when appropriate: carbenicillin (200 µg/mL), gentamicin (30 µg/mL), tetracycline (100 µg/mL). *Escherichia coli* strains for cloning were cultured in lysogeny broth (LB; 1% tryptone, 0.5% yeast extract, 1% sodium chloride). The following antibiotics were added for *E. coli* when appropriate: ampicillin (100 µg/mL), gentamicin (15 µg/mL), tetracycline (12 µg/mL). All strains used in this study can be found in the Key Resources Table.

### Generating *P. aeruginosa* mutants

*P. aeruginosa* mutants in this study were generated by allelic exchange at the native site in the chromosome using Gibson Assembly with a pEXG2-Tc vector containing the DNA mutation between restriction sites SacI and XbaI (Hmelo et al., 2015). Mutant constructs were generated by PCR amplifying ∼1 kb DNA fragments upstream and downstream of the gene or region of interest while substitution mutants were generated by synthesis of a DNA fragment gene block containing the correct codon change (Integrated DNA Technologies, Coralville, IA, USA). Assembled vectors were transformed into *E. coli* DH5*α*, then into *E. coli* S17 for conjugation into *P. aeruginosa*. Correct mutations in *P. aeruginosa* were verified with PCR and Sanger sequencing. *P. aeruginosa ΔpilK*, *ΔchpB*, *ΔpilJ* deletion mutants and *pilJ* LBDs and methylation site mutants were complemented by electroporating the respective mutant with expression vector pMQ80 containing the full-length gene under control of the arabinose-inducible PBAD promoter and fused to a C-terminal GFP tag(Shanks et al., 2006). All oligonucleotides used to generate *P. aeruginosa* mutants can be found in Table 1.

**Table 1.**
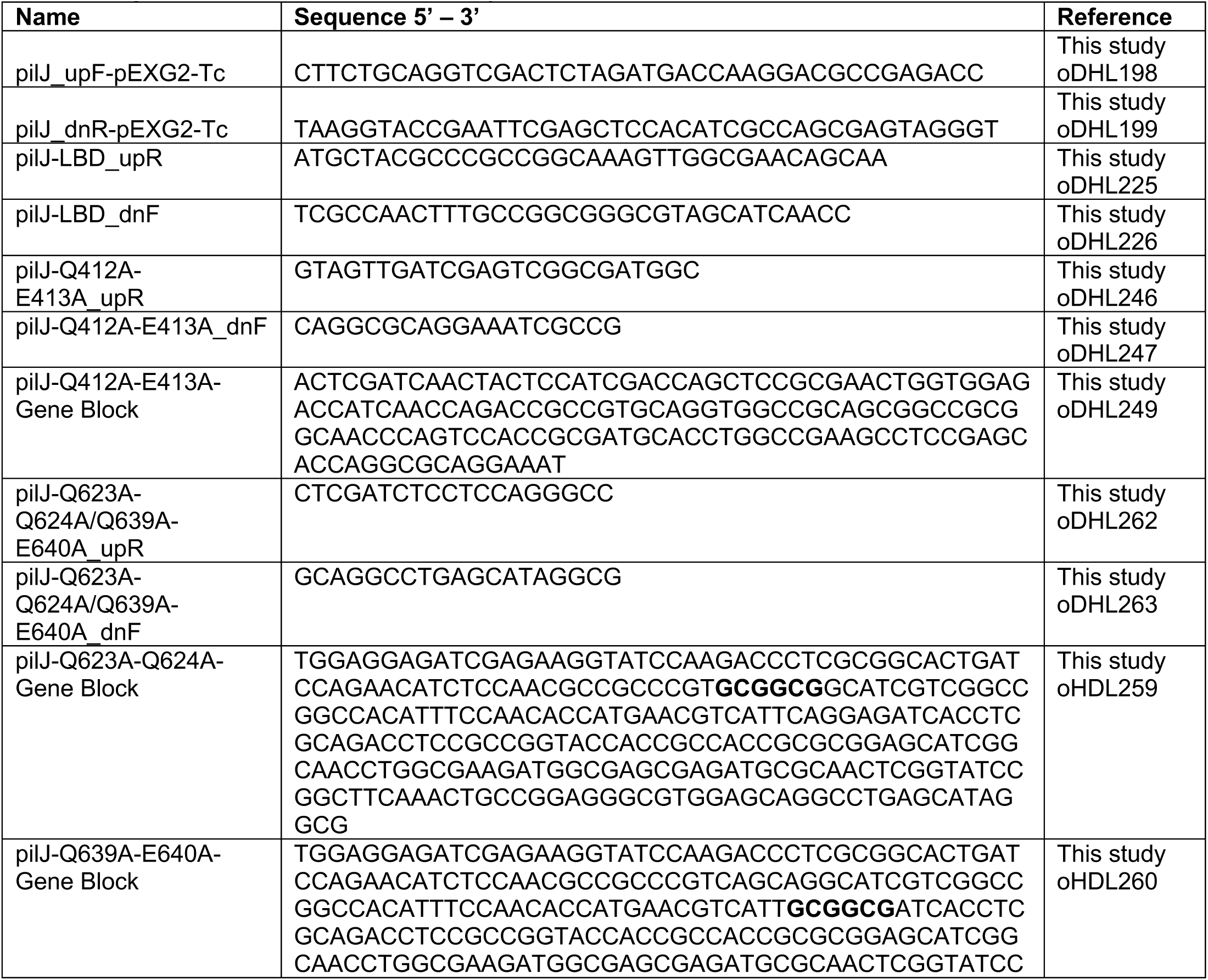
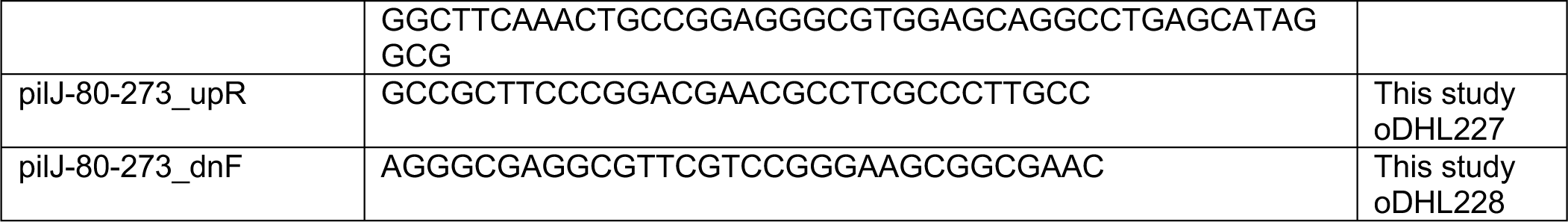
Oligonucleotides used in this study.

### Macroscopic coculture twitching chemotaxis assay

Motility experiments were performed as previously described (Kearns et al., 2001; Limoli et al., 2019; Miller et al., 2008). Buffered agar plates (recipe: 10 mM Tris, pH 7.6; 8 mM MgSO4; 1 mM NaPO4, pH 7.6; and 1.5% agar) were poured and allowed to solidify for 1 hour prior to incubation for 16 hours at 37°C and 22% humidity. After solidifying, 4 µL of either growth medium (TSB) or cell-free supernatant derived from an overnight culture of *S. aureus* at OD_600_ 5.0 and filter sterilized with a 0.22 µm filter were spotted on the surface of the plate and allowed to diffuse for 24 hours at 37°C and 22% humidity to establish a gradient. *P. aeruginosa* cultures were incubated overnight in TSB with aeration at 37°C, subcultured 1:100 in TSB the following morning, then standardized to OD_600_ 12.0 in 100 µL of 1 mM MOPS buffer supplemented with 8 mM MgSO_4_ prior to inoculating 1 µL on the surface of the plate at five mm from the center of the gradient. Plates were incubated in a single layer, agar-side down, for 24 hours at 37°C with 22% humidity, followed by an additional 16 hours at room temperature prior to imaging the motility response of *P. aeruginosa.* Images were captured using a Zeiss stereoscope with Zeiss Axiocam 506 camera and directional motility ratios were calculated in Fiji before graphing and performing statistical analysis in GraphPad Prism.

### Macroscopic coculture subsurface twitching assay

The assay was modified from a previous protocol, as shown before (Limoli et al., 2019; Turnbull & Whitchurch, 2014). Prior to pouring plates, 200 µL of either growth medium (TSB) or cell-free supernatant derived from an overnight culture of *S. aureus* at OD_600_ 2.0 and filter sterilized with a 0.22 µm filter were spread on the bottom of a 120 mm square petri plate. Tryptic soy agar (1.5%) was then poured into the plates and allowed to dry for 4.5 hours at room temperature. *P. aeruginosa* cultures were incubated overnight in TSB with aeration at 37°C, subcultured 1:100 in TSB the following morning, then standardized to OD_600_ 2.0 in 1 mL TSB. A sterile toothpick was dipped into the standardized culture, then stabbed to the bottom of the agar plate. Plates were incubated in a single-layer, agar-side down, at 37°C with 22% humidity for 24 hours, followed by an additional 24 hours at room temperature. Following incubation, the agar was removed from the plates and the motility diameters were measured in mm. Diameter measurements were graphed and analyzed in GraphPad Prism.

### Live-imaging and tracking of *P. aeruginosa* directional response to *S. aureus*

To visualize single-cell motility behavior and measure cAMP in individual cells, *P. aeruginosa* cells were imaged using a previously described method (Limoli et al., 2019; Yarrington et al., 2020). *P. aeruginosa* and *S. aureus* were grown in M8T medium overnight at 37°C with aeration, subcultured the next day in fresh M8T and grown to mid-log phase at 37°C with aeration. Cultures were standardized to OD_600_ of 0.015 to 0.03 for *P. aeruginosa* or 0.05 to 0.1 for *S. aureus*. For coculture experiments, *P. aeruginosa* and *S. aureus* were mixed 1:1. One µL of mono- or coculture cells were inoculated onto a 10 mm diameter glass coverslip in a 35 mm dish before placing an agarose pad on top. Pads were made by pipetting 920 µL of M8T with 2% molten agarose into a 35 mm dish containing a 10 mm diameter mold and drying uncovered for 1 hour at room temperature, followed by 1 hour 15 minutes at room temperature covered with a lid, then 1 hour at 37°C before transferring the pad onto the inoculated coverslip. Time-lapse imaging was performed with an inverted Nikon Ti2 Eclipse microscope, 100x oil objective (1.45 NA), and Andor Sona camera. Phase contrast images were acquired every 20 minutes for 2 hours, then every 1 second for 3-4 hours with 100 ms exposure and 20% DIA LED light. Fluorescent images were acquired every 20 minutes with TxRed images taken at 100 ms exposure and 20% Sola fluorescent light and YFP imaged at 25 ms exposure and 20% Sola fluorescent light.

Images were analyzed using Nikon NIS-Elements AR software. To automatically track movements of single cells, bacterial cells in the phase channel were first converted into binary objects by thresholding to the dark bacterial cells. The Tracking Module in NIS-Elements was then used to form trajectories for all binary objects. Specifically, the following parameters were used: objects have a minimum area of 1 µm^2^, track with random motion model, find center of object based on area, no maximum speed limit of each object track, allow new tracks after the first frame in file and track objects backwards to previous frames. Tracks were allowed up to 60 gaps (frames) per track and any tracks with less than 180 frames were automatically removed from the final trajectories. Trajectories were exported and analyzed in Python using the parameters described below.

### Principal direction of single-cell trajectories

The gyration tensor, commonly employed in polymer physics determines the principal direction of each single-cell trajectory *i* (Kim & Baig, 2016). The shape of each random walk is described by the gyration tensor

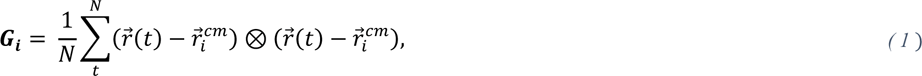

for a trajectory of *N* time steps each at position *r⃗*(*t*) at time *t* and center of mass position 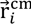 Eigendecomposition of the gyration tensor determines the largest eigenvalue and the associated eigenvector 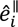 is identified as the principal direction of motion for the *i*^th^ trajectory. The sign of 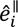 is chosen by setting the direction of motion parallel to the end-to-end vector. The principal direction is determined without any information about the location of *S*. *aureus* colonies or other *P. aeruginosa* cells. Rose graphs shown in Figures 2-4 present histograms of 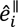.

### Directed Mean-Squared Displacement

The mean-squared displacement (MSD)

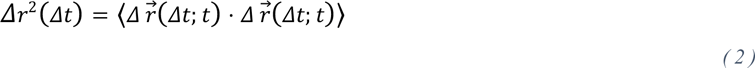

of displacement vectors *Δr⃗*(Δ*t*; *t*) = *r⃗*(*t* + *Δt*) − *r⃗*(*t*) is a measure of the distance gone after a lag time *Δt*, where the average 〈⋅〉 is the ensemble averaover all trajectories *i* and all times *t* within that trajectory. While the MSD quantifies the degree of motion, it assumes isotropic dynamics and so does not discern between any potential directionality in microbial motion. Quantifying the degree of motion parallel and perpendicular to the principal direction requires decomposing the movement into components parallel and perpendicular to each principal direction 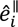. The displacement in the principal direction is 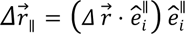and the orthogonal direction is *Δr⃗*_⊥_ = *Δr⃗* − *Δr⃗*_∥_. From these, the parallel and perpendicular MSD are

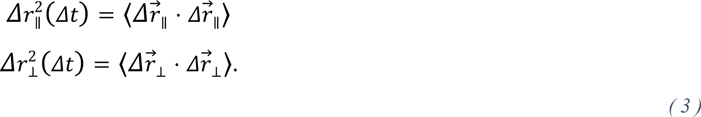

To characterize the MSDs, the anomalous exponent α defined by the power law

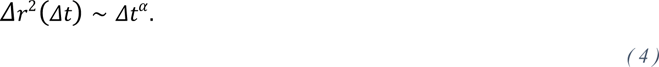

Exponents of α = 1 describe diffusive dynamics, while α < 1 represents subdiffusive motion and α > 1 superdiffusive.

### Displacement probability distributions

The MSD can be deceiving since it is known that a broad class of diffusive dynamics exist in soft matter, biological, and complex systems for which the dynamics are “Brownian yet non-Gaussian” (Chechkin et al., 2017; Metzler, 2020). In such systems, the MSD appears diffusive with anomalous exponent α = 1 but the probability density function (PDF) of steps *P*(*Δr*; *Δt*) is non-Gaussian, a traditional assumption for Brownian motion. The probability of finding a bacterium *Δr* after some lag time *Δt* is called the van Hove self-correlation function and it has proven useful in understanding the dynamics of simulations of twitching bacteria (Nagel et al., 2020). The formal definition of the van Hove self-correlation function is

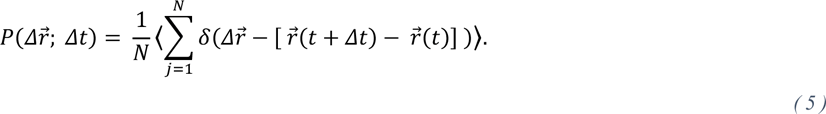

To consider the probability that *P. aeruginosa* cells move a given distance towards *S. aureus* colonies, the van Hove function is found for steps parallel or orthogonal to the principal direction of motion, *Δr⃗*_∥_ and *Δr⃗*_⊥_ respectively. There are two probability density functions of particular interest here.

*(i)* The first is a Gaussian diffusive distribution

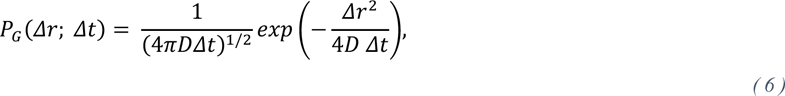

for a random process with diffusion coefficient *D*. This form can be scaled in time to collapse the distribution to 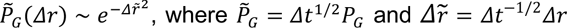. Thus, observing a Gaussian distribution at only one lag time is insufficient for determining Fickian diffusivity. An example of this is the microscopy imaging noise (Figure 2 – figure supplement 4A): although the step size distributions are Gaussian, they do not scale in time for lag times *Δt* < 300*s*, a distinct indication that this is a measure of the tracking noise and not diffusion of the dust particle.
*(ii)* The second probability density functions of interest is a Laplace distribution

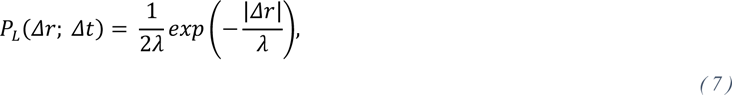

for a decay length *λ*. Laplace distributions have longer tails than Gaussian distributions and have emerged as a canonical example of non-Gaussian functions that lead to Brownian MSDs (Chechkin et al., 2017; Metzler, 2020). If the decay length scales with lag time as *λ* = (〈*D*〉 *Δt*)^1/2^ for an average diffusivity 〈*D*〉 then the MSD scales diffusively. In this case, the distributions can be collapsed with the same scaling as the Gaussian distributions, 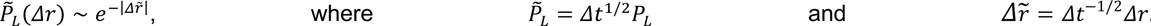.

The probability density distributions of step sizes are typically dominated by a sharp narrow peak of highly-likely small step sizes, which represents jiggling and long tails of rare-but-large step sizes (Figures 2-4) (Kühn et al., 2021). The long tails are primarily exponential. It is tempting to think that the narrow peak of small steps sizes represents the imaging uncertainty. However, we find that the standard deviation of the imaging distribution is 1.4 × 10^-2^*μm* (Figure 2 – figure supplement 4A), narrower than the width of the primary van Hove peak (Figure 2 – figure supplement 4B). Indeed, the narrow peak is not Gaussian at all, but rather better fit by a second Laplace distribution. Thus, the displacement probability density distributions are well described as double Laplace functions (Figure 2 – figure supplement 4B).

### Identifying subpopulations of persistent movers and resters

Qualitative assessment of the microscopy data makes it apparent that *P. aeruginosa* cells possess two dynamic modes:

1. Persistent “Resters”: Colony-associate *P. aeruginosa* cells do not exhibit an active exploratory motion. Instead, the motion of these cells is composed of small ‘jiggling’ and expansion due to colony growth.
2. Persistent “Movers”: These are cells that have left the colony to actively move through the surroundings, either as individuals or in multi-cell rafts. Like “resters”, these “movers” exhibit small jittering motion but also intermittently persistent motion. The intermittency of the motion can at times have a run-reversal-type or a run-rest-type character, but a “mover” is not simply a continually moving bacterium.

Both of these dynamic modes are composed of small jittery motion and larger motions, which makes it difficult to algorithmically separate the cells into subpopulations of movers and resters.

To disentangle the subpopulations, we consider the velocity-velocity correlation function 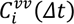 for lagtime *Δt* for each bacterium *i* averaged over start times. The velocity autocorrelation function takes into account both the direction and speed of the bacteria. Rather than averaging the correlation function over all possible start times within trajectory *i*, we employ a rolling velocity autocorrelation

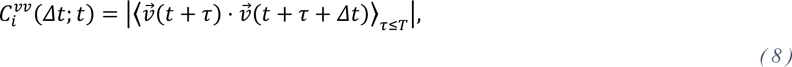

where *t* is each time point in the trajectory, *T* is the rolling window duration, τ is every possible starting time within the rolling window and the average ⟨⋅⟩_*r*≤*T*_ is over all starting times. Since run-reversal dynamics are likely to be an aspect of the dynamics, we consider the absolute value of the velocity autocorrelation. If the duration *T* is too short then the correlation functions are overly noisy but if it is too long then instances of correlated motion is smeared out. To assess the immediate degree of correlation in motion, the correlation function is averaged over the duration to produce a correlation constant 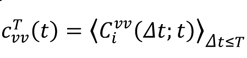. The correlation constant acts as a signal of immediately persistent motion, with persistent resters showing near-zero 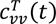 and movers having significantly larger values above a cutoff *c*^∗^. While 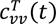 matches our expectations from qualitative observations of the microscopy movies, false positive instances of colony-associated cells occur. Thus, the signal is weighted by the behavior of neighboring cells

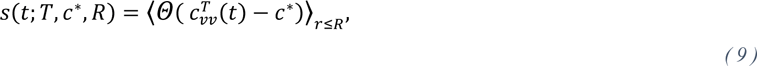

where *Θ*(⋅) is the Heaviside step function and the average is over all neighbouring cells in the vicinity of *r* ≤ *R*. Finally, the neighbor-weighted signal *s* is given a cutoff *s*^∗^, above which cells are identified as “persistent movers” and below which they are “persistent resters.” The parameters are chosen to be *T* = 30*s*, *c*^∗^ = 0.01, *R* = 6*μm* and *s*^∗^ = 0.75 for this study. Due to the intermittent nature of the twitching dynamics, once a bacterium has been identified as a mover, it keeps a mover-designation until the trajectory is lost.

### Quantification of intracellular cAMP

To quantify cAMP in individual cells, time-lapse imaging was performed with *P. aeruginosa* cells carrying the P*xphA-yfp* P*rpoD-mKate2* dual fluorescent reporter(Persat et al., 2015). Thresholding of bacterial cells in the red channel to the constitutively expressed P*rpoD*-*mKate2* fluorescence generated binary objects. The fluorescence of these binaries was then measured in the YFP channel for levels of P*xphA*-*yfp* expression. For total cAMP in a frame at a given time point, the ratio of YFP over mKate2 intensity for each bacterial cell was calculated, then summed with all other bacterial cells. For normalization, the total YFP/mKate2 ratio from all objects in the frame was normalized to the average area of all binary objects in the same frame. The ratios for each time point were then graphed and analyzed in GraphPad Prism.

## Acknowledgements

This work was supported by funding from the CFF Postdoc-to-Faculty Transition Award (LIMOLI18F5), CFF RDP Junior Faculty Recruitment Award (LIMOLI19R3), NIH (R35GM142760), CFF Student Traineeship Award (YARRIN21H0), and the European Research Council (ERC) under the European Union’s Horizon 2020 research and innovation programme (Grant agreement No. 851196). We thank Drs. George O’Toole and Sherry Kuchma for *P. aeruginosa* and *E. coli* bacterial strains. We also thank Dr. J. Muse Davis for use of the stereoscope and Dr. Timothy Yahr for the pEXG2-Tc cloning vector. We are grateful to George O’Toole and members of the Limoli Lab for thoughtful discussions and feedback on the manuscript.

---

**Figure 2 – Figure supplement 1.**
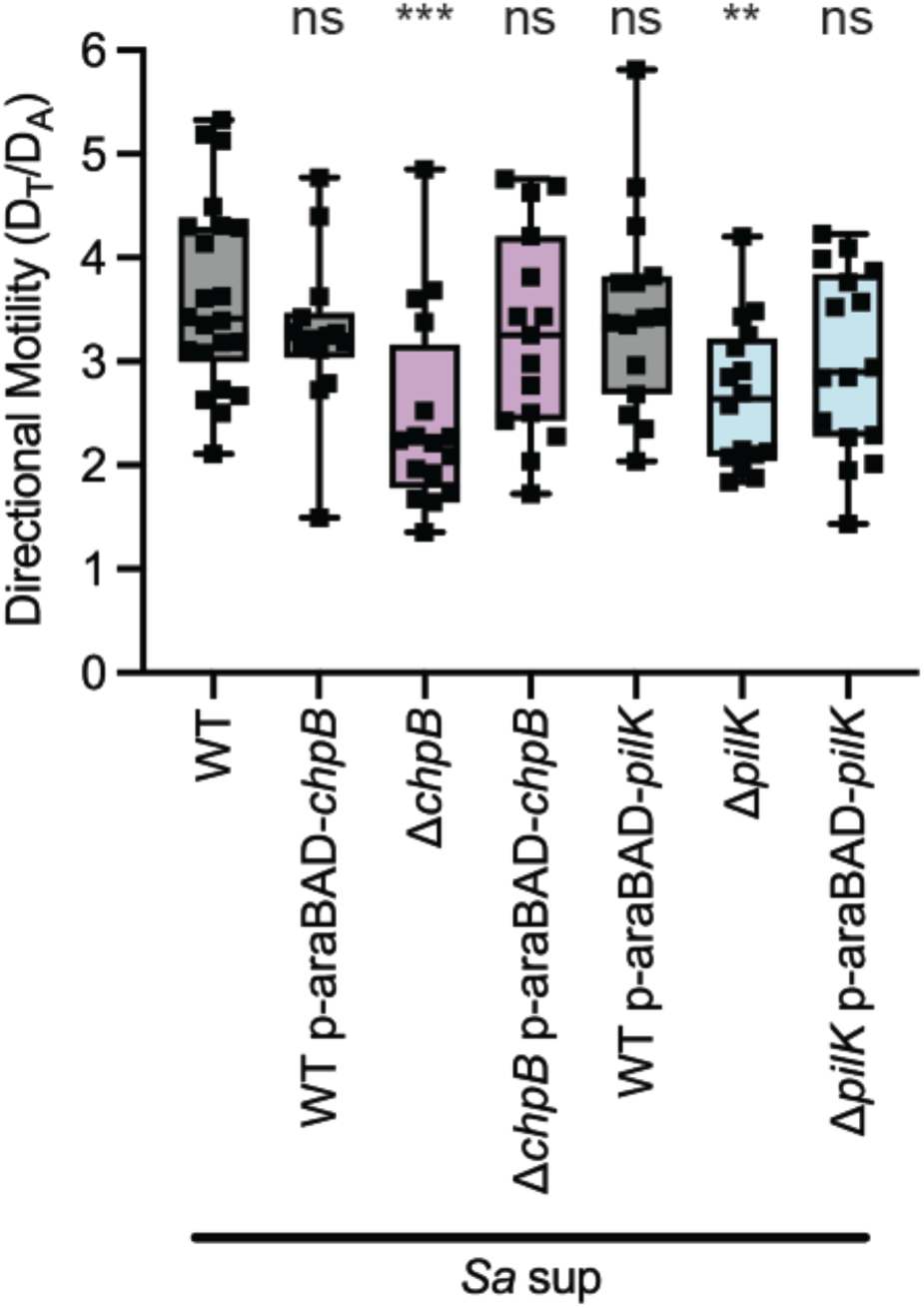
Complementing Δ*pilK,* and Δ*chpB* restores pilus-mediated chemotaxis. Directional motility towards *S. aureus* secreted factors of wildtype*, ΔpilK*, and *ΔchpB* with and without complementing plasmids carrying arabinose-inducible copies of *pilK* or *chpB*. Complemented strains were induced with 0.2% arabinose; however, phenotypes were the same in the absence of induction. Directional motility for at least four biological replicates, each containing a minimum of three technical replicates are shown. Statistical significance was determined with a one-way ANOVA followed by Dunnett’s multiple comparisons test. *\*\*\** indicates p< 0.001; ** indicates p<0.01; *ns* indicates no statistically significant difference.

**Figure 2 – Figure supplement 2.**
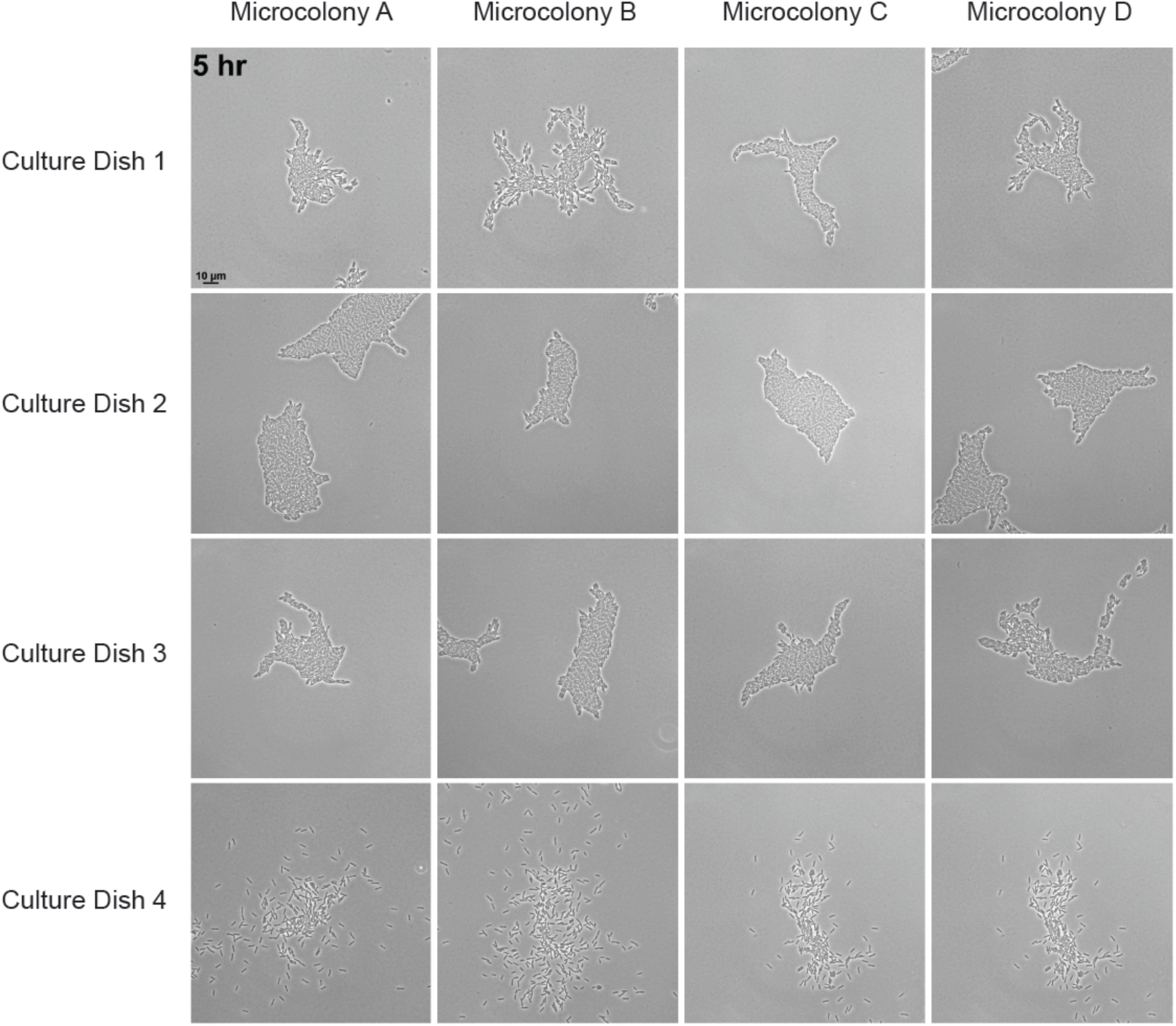
*P. aeruginosa* Δ*pilK* has bimodal pilus-mediated motility. Separate *ΔpilK* cultures plated onto four individual experimental dishes and four fields of view in each culture dish were simultaneously imaged at 5 hours post-inoculation. Agarose pads were made from the same media and dried under the same conditions at the same time. A range in motility phenotypes are seen between all microcolonies imaged.

**Figure 2 – figure supplement 3.**
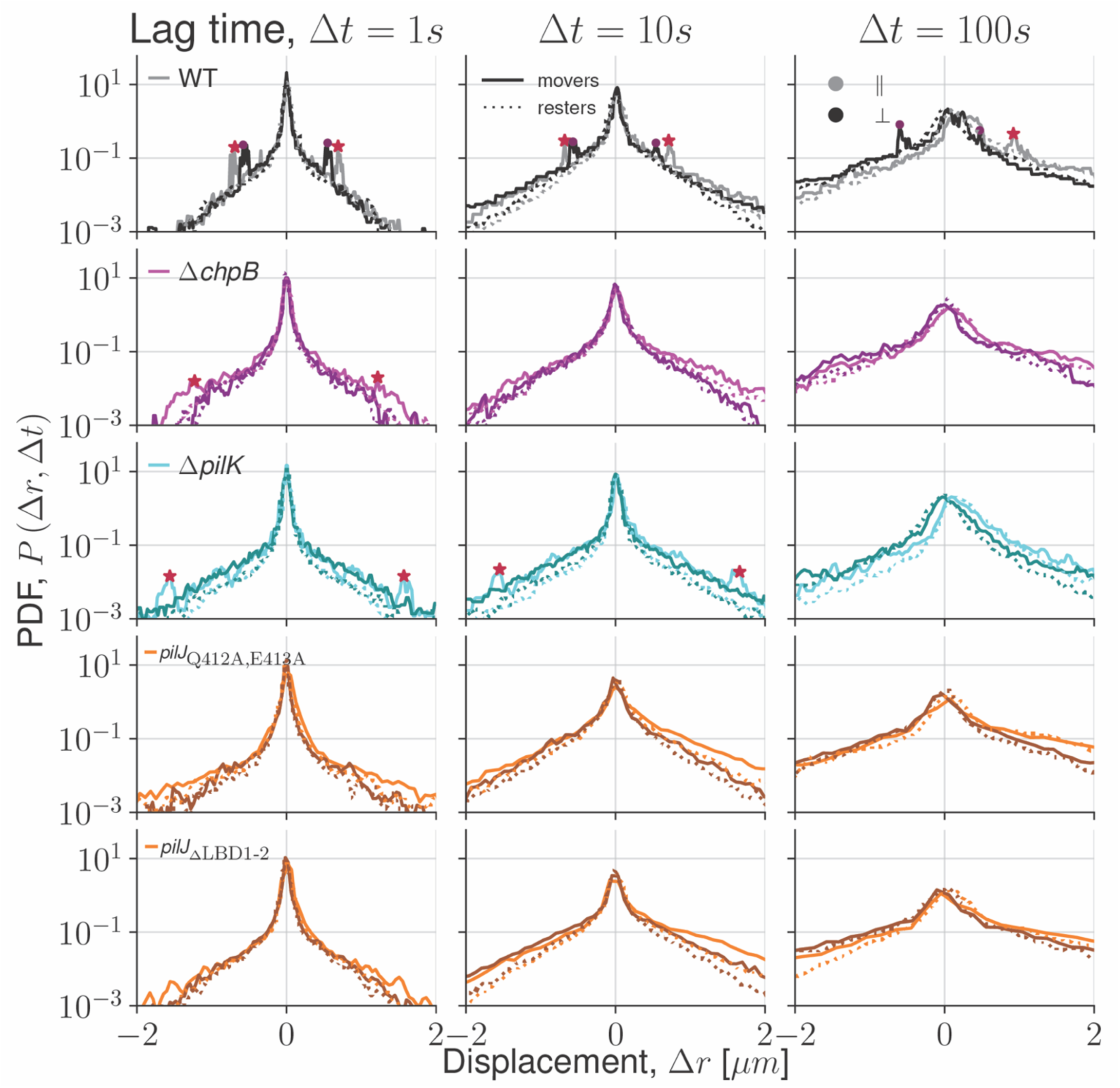
Van Hove distributions for all mutants. Step size distributions for each *P. aeruginosa* strain are displayed by distance of displacement (*Δr*) of cells. Step size probability density functions (PDF) are shown for movers (solid lines) and resters (dotted lines) in the parallel (lighter lines, ||) and perpendicular (darker lines, ⊥) directions. Step sizes for each *P. aeruginosa* strain were calculated from cell trajectories with a 1 second (left), 10 second (middle), and 100 second (right) lag time (*Δt*). Red stars (movers) and maroon dots (resters) highlight the non-zero sharp-shoulder peak step size, when present.

**Figure 2 – figure supplement 4.**
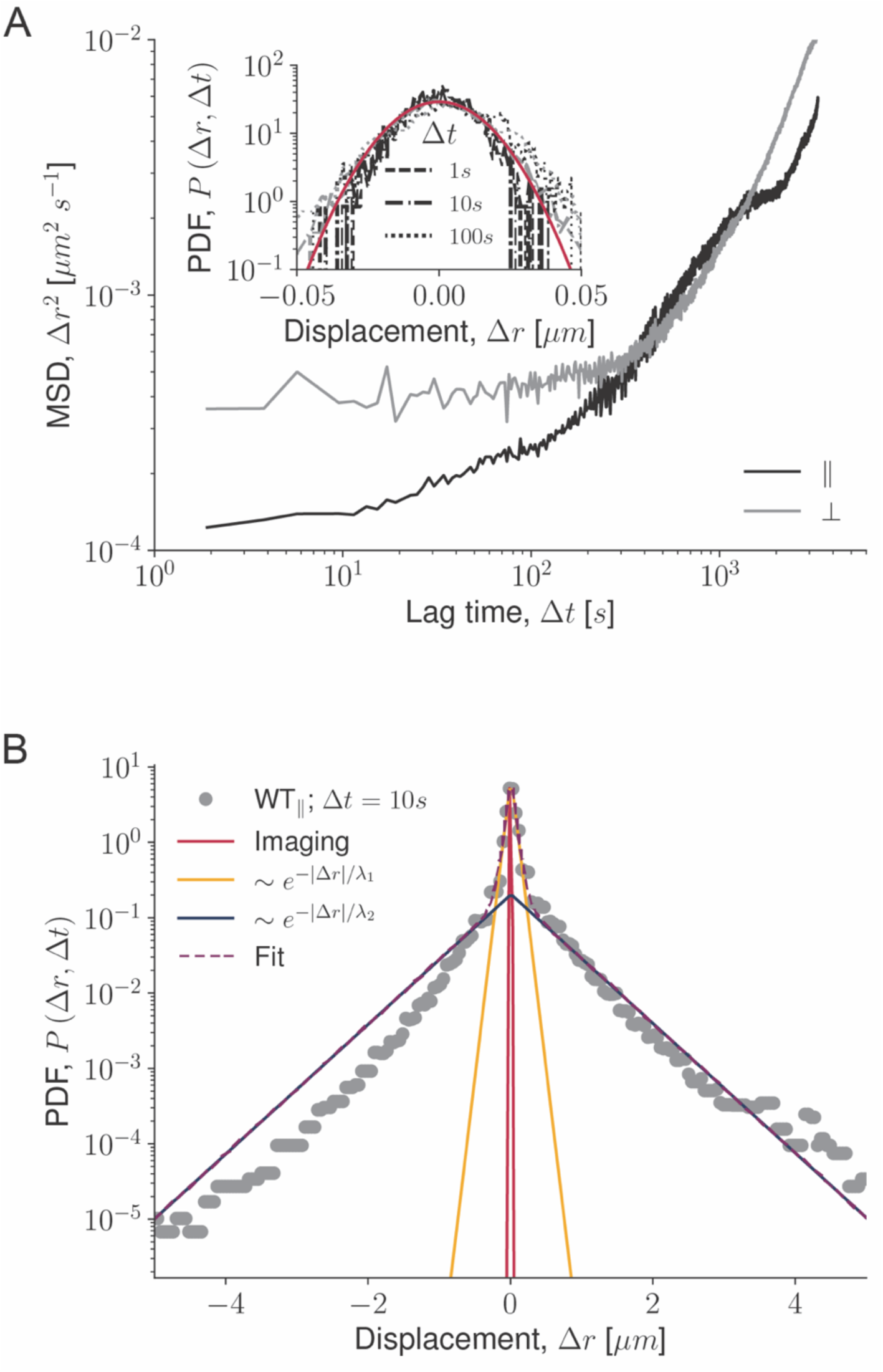
Dynamics due to imaging errors. (A) Mean-squared displacements (MSD) for the parallel (||) and perpendicular (⊥) directions for dust particles used to measure noise in the imaging. Inset shows the particle-displacement probability density function (PDF). The PDF is a narrow noise peak, that is fit to a Gaussian distribution (solid red line) but is non-diffusive, as it does not broaden in time. (B) PDF of the total cell step displacements (*Δr*), regardless of principal direction, for wildtype cells at a lag time of *Δt=10* seconds. The PDF is composed of a narrow peak of small displacements (jiggling) and long tails of large-but-rare displacements. The narrow peak cannot be explained by imaging uncertainty (solid red curve) and is better described by a Laplace distribution (Eq 7), as are the long tails.

**Figure 3 – Figure supplement 1.**
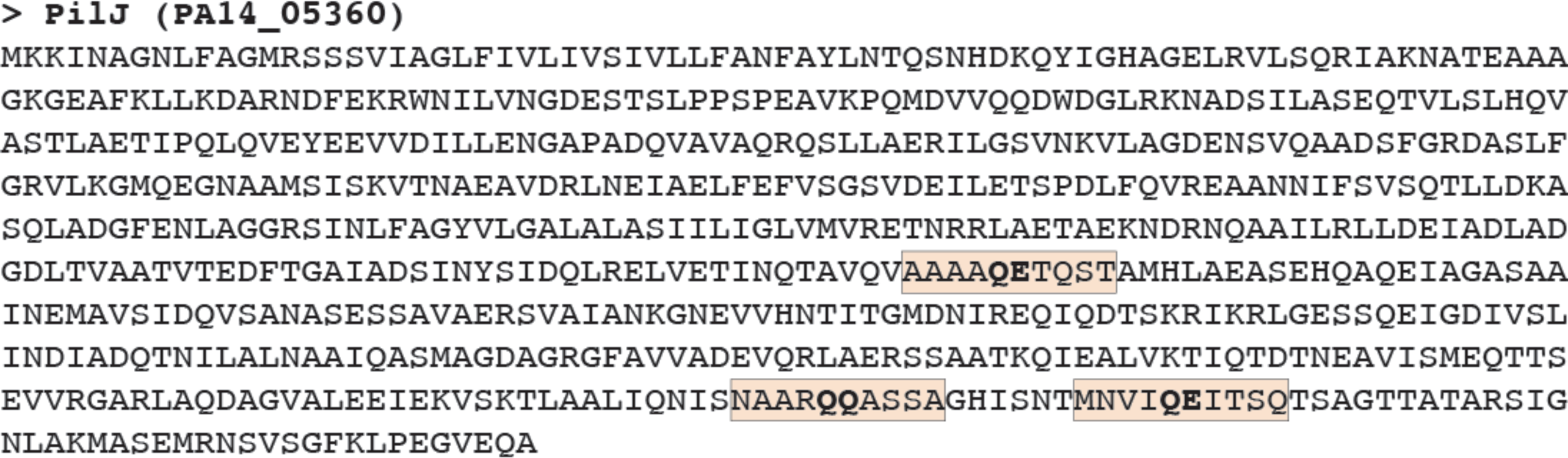
Methylation sites of PilJ. Amino acid sequence of *P. aeruginosa* PA14 PilJ (PA14_05360) with conserved MCP methylation motifs highlighted in pale orange and predicted methyl modification glutamate/glutamine residue pairs in bold.

**Figure 3 – Figure supplement 2.**
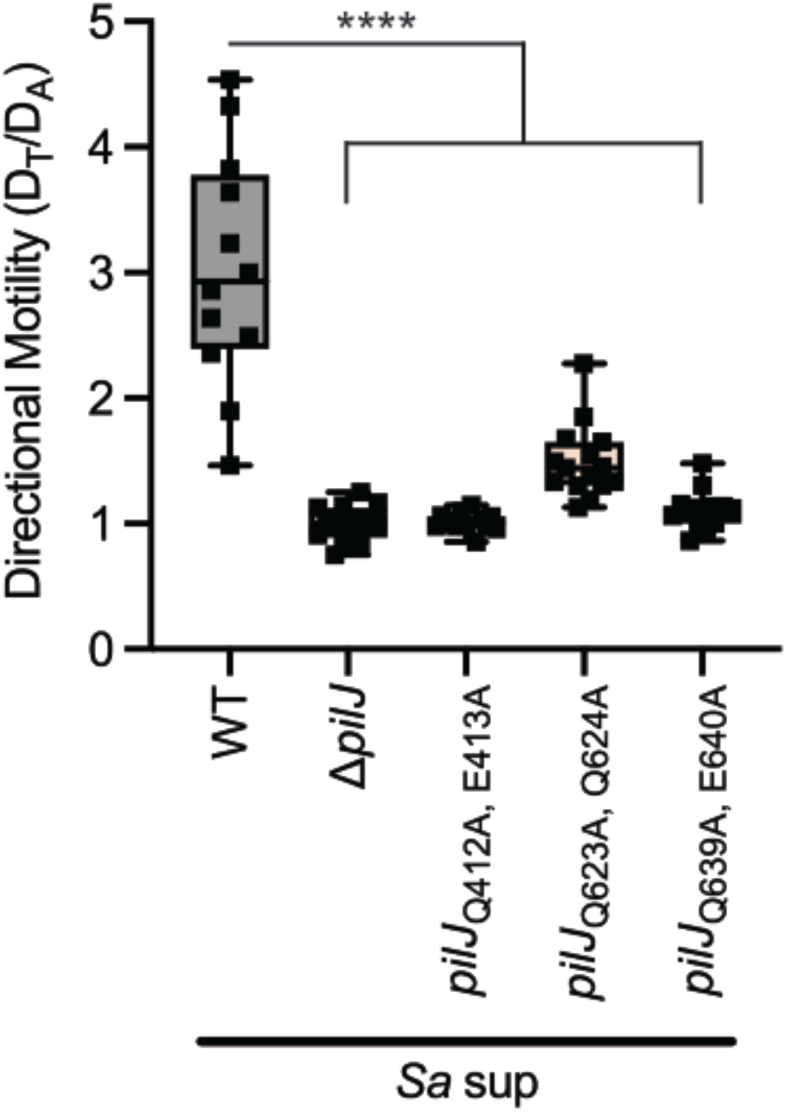
Mutation of any cytoplasmic PilJ methylation site reduces *P. aeruginosa* attraction to interspecies signals. Directional motility of *P. aeruginosa* wildtype and methylation mutants *pilJ*Q_412A, E413A_, *pilJ*Q_623A, Q624A_, and *pilJ*Q_639A, E640A_ in the presence of a gradient of *S. aureus* supernatant. Directional motility for at least four biological replicates, each containing a minimum of three technical replicates are shown and statistical significance was determined with a one-way ANOVA followed by Dunnett’s multiple comparisons test. **** indicates p<0.0001.

**Figure 3 – Figure supplement 3.**
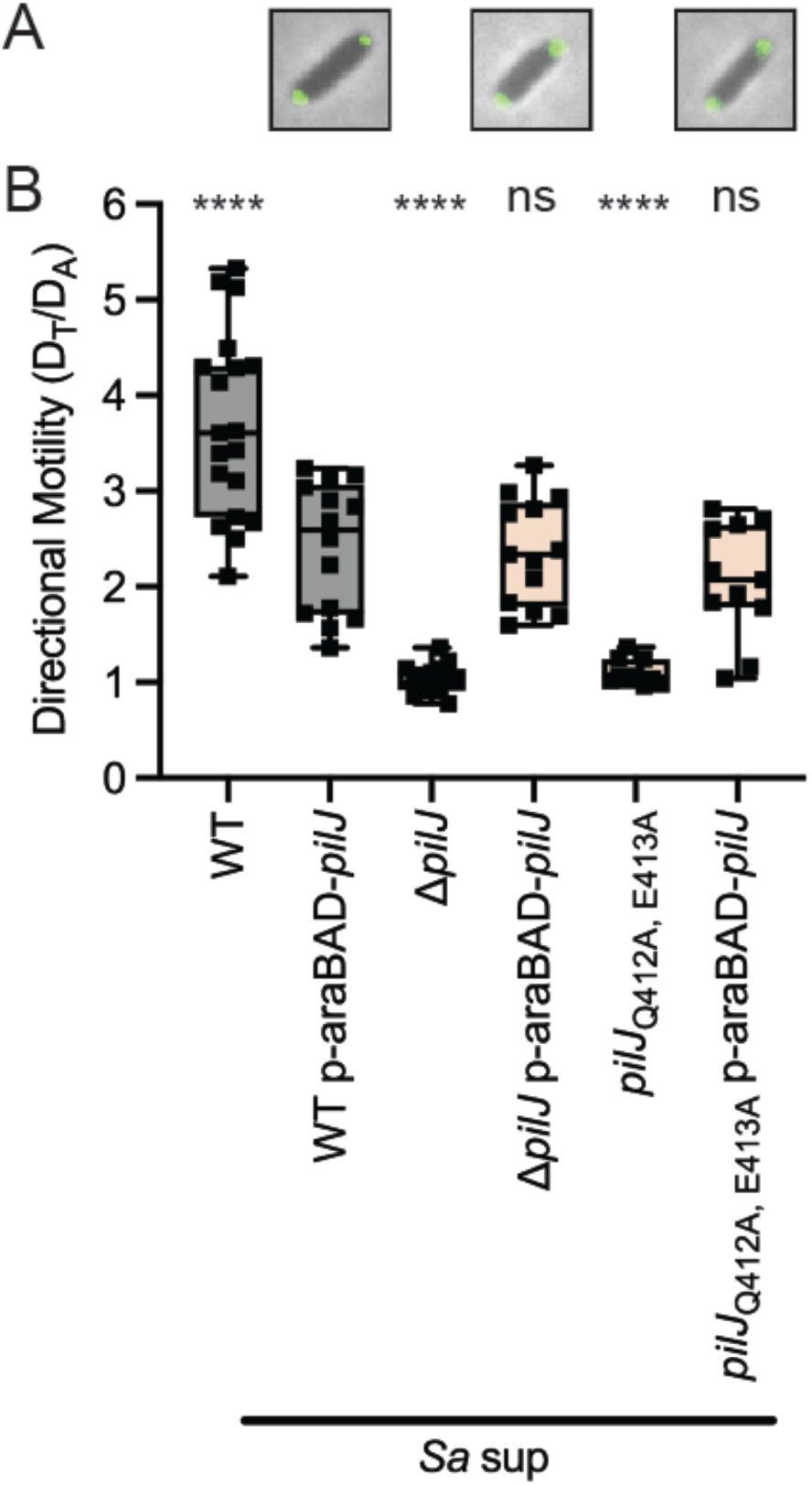
Complemented *pilJ*Q_412A, E413A_ with full-length PilJ displays bipolar PilJ localization and restoration of TFP-mediated directional motility. (A) Representative *P. aeruginosa* cells with bipolarly localized GFP-tagged PilJ. (B) Directional motility towards *S. aureus* secreted factors of wildtype or *pilJ*Q_412A, E413A_ with and without complementing plasmids carrying arabinose-inducible copy of wildtype *pilJ*. Complemented strains were induced with 0.2% arabinose; however, phenotypes were the same in the absence of induction. Directional motility for at least three biological replicates, each containing a minimum of three technical replicates are shown and statistical significance was determined with a one-way ANOVA followed by Dunnett’s multiple comparisons test to compare each strain to wildtype *P. aeruginosa* carrying p-araBAD-*pilJ*. **** indicates p<0.0001; *ns* indicates no statistically significant difference.

**Figure 4 – Figure supplement 1.**
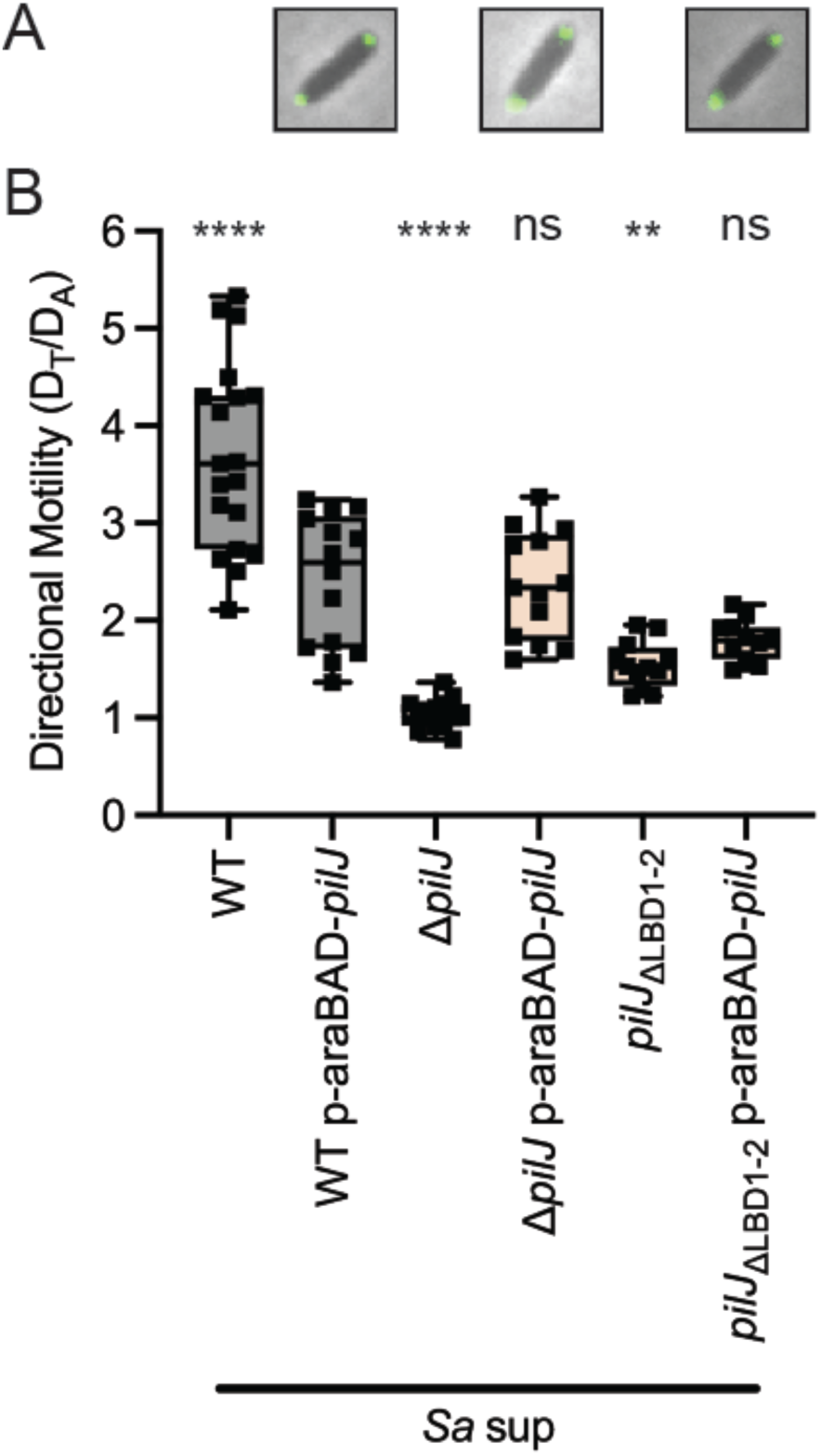
Complemented *pilJ*_ΔLBD1-2_ has bipolarly localized PilJ but only partial restoration of TFP-mediated chemotaxis. (A) Representative *P. aeruginosa* cells with bipolarly localized GFP-tagged PilJ. (B) Directional motility towards *S. aureus* secreted factors of wildtype or *pilJ*_ΔLBD1-2_ with and without complementing plasmid carrying arabinose-inducible copy of wildtype *pilJ*. Complemented strains were induced with 0.2% arabinose; however, phenotypes were the same in the absence of induction. Directional motility for at least three biological replicates, each containing a minimum of three technical replicates are shown and statistical significance was determined with a one-way ANOVA followed by Dunnett’s multiple comparisons test to compare each strain to wildtype *P. aeruginosa* carrying p-araBAD-*pilJ*. **** indicates p<0.0001; ** indicates p<0.01; *ns* indicates no statistically significant difference.

**Figure 4 – figure supplement 2.**
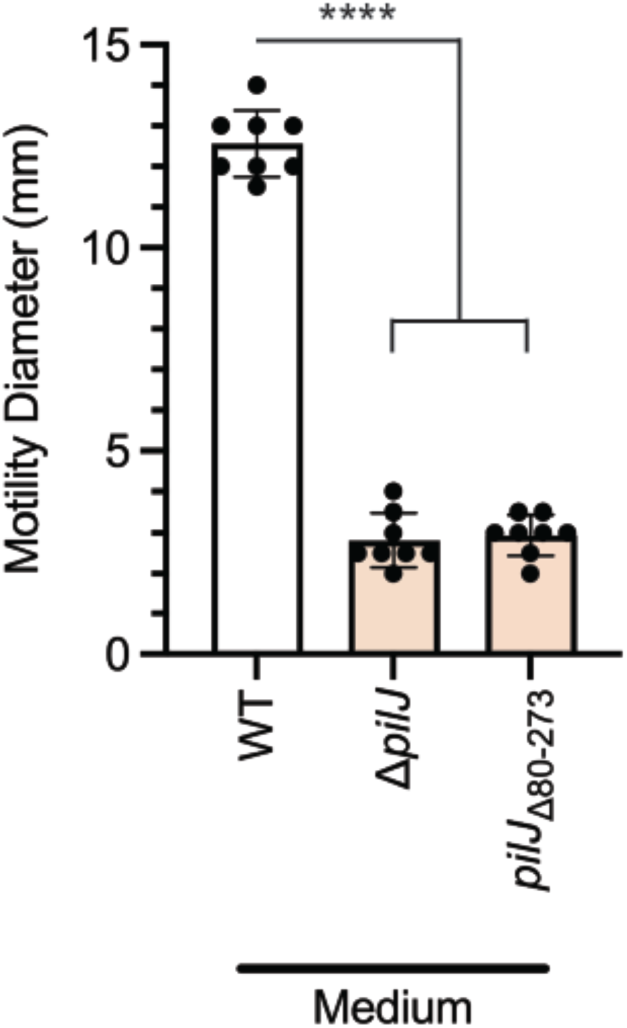
PilJ mutant lacking a portion of the periplasmic region is non-motile. Twitching motility diameters of *P. aeruginosa* wildtype, *ΔpilJ*, and a *pilJ* mutant lacking amino acids 80-273 (*pilJ*Δ80-273). Macroscopic motility measurements are shown for two biological replicates, each containing four technical replicates and statistical significance was determined with a one-way ANOVA followed by Dunnett’s multiple comparisons test. *\*\*\*\** indicates p< 0.0001.

**Figure 5 – figure supplement 1.**
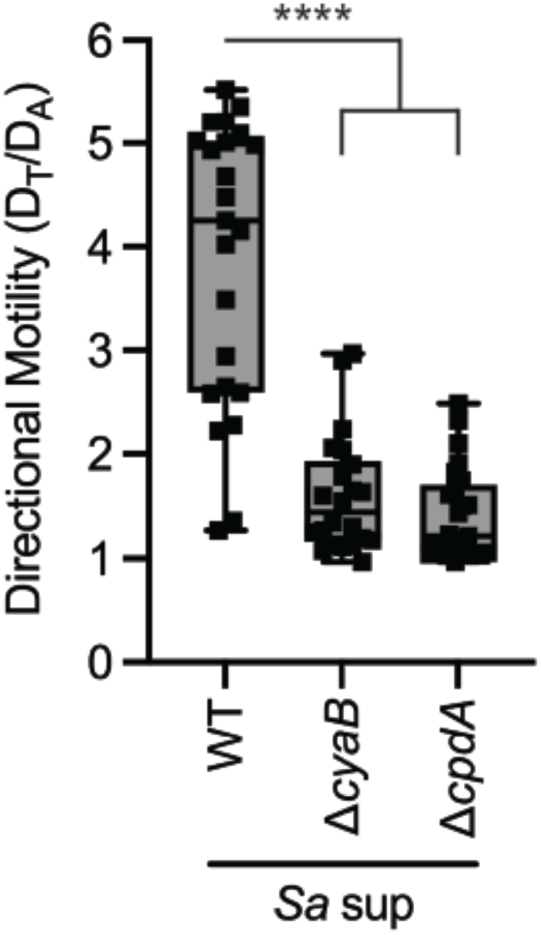
Enzymes that control cAMP levels are necessary for pilus-mediated chemotaxis. Directional motility towards *S. aureus* secreted factors of *ΔcyaB* and *ΔcpdA*. At least three biological replicates, each containing a minimum of three technical replicates are shown and statistical significance was determined with a one-way ANOVA followed by Dunnett’s multiple comparisons test. *\*\*\*\** indicates p< 0.0001.

